# *Caenorhabditis elegans* Uses Canonical and Non-canonical Hippo signaling

**DOI:** 10.1101/2025.08.22.671798

**Authors:** Linh Huynh, Razan A. Fakieh, C’Brionne Hendrix, Reid Powell, David J. Reiner

## Abstract

Hippo signaling is a conserved regulator of tissue homeostasis across metazoans. The Ste20 family kinase Hippo/MST activates the NDR family kinase Warts/LATS to inhibit the transcriptional coactivator Yorkie/YAP/TAZ and its transcription factor partner Scalloped/TEAD. In *Caenorhabditis elegans*, cell lineages and organ sizes are largely invariant, and classical Hippo phenotypes such as tissue overgrowth are absent. Nevertheless, WTS-1, YAP-1, and the TEAD-like transcription factor EGL-44 form a conserved core module required for larval development past the L2 stage. Crucially, a direct role for Hippo signaling remains unestablished. To address this question, we generated a fluorescently tagged endogenous YAP-1 as a live biomarker of pathway activity. Upon WTS-1 loss, endogenous YAP-1 translocated from cytosol to nucleus in epithelium and intestine. Tissue-specific depletion revealed that intestinal, but not epithelial, WTS-1 is essential for progression past L2. The duplicated Hippo-related kinases CST-1 and CST-2 repressed YAP-1 nuclear localization in the epithelium but not intestine, indicating that intestinal WTS-1 functions without CST-1/2. The Ste20 kinase MIG-15, orthologous to *Drosophila* Misshapen and mammalian MAP4K4/6/7/8, was redundant with CST-1/2 for larval progression. Yet deficient MIG-15 uniquely increased YAP-1 abundance without driving nuclear localization. By contrast, the Ste20 kinase GCK-2, orthologous to *Drosophila* Happyhour and mammalian MAP4K1/2/3/5, had no detectable role. Our findings establish *C. elegans* as a model for Hippo signaling, with a canonical cascade active in the epithelium and noncanonical inputs controlling WTS-1 in the intestine. In this context, YAP-1/EGL-44 outputs are repurposed from growth control to non-proliferative developmental functions.

## INTRODUCTION

The Hippo signaling pathway was first discovered in *Drosophila* loss-of-function screens that identified tumor suppressor genes required to prevent tissue overgrowth (JUSTICE *et al*. 1995; TAPON *et al*. 2002; HARVEY *et al*. 2003; WU *et al*. 2003). Subsequent studies revealed that the downstream transcriptional coactivator Yorkie (Yki) is essential to avoid the converse phenotype of tissue undergrowth (HUANG *et al*. 2005). The canonical pathway consists of the Ste20-family (from yeast <u>Ste</u>rile <u>20</u>) kinase Hippo (Hpo; mammalian Mst1/2) phosphorylating and activating the NDR-family kinase Warts (Wts; mammalian LATS1/2), which in turn phosphorylates and represses Yki/YAP/TAZ (HUANG *et al*. 2005; OH AND IRVINE 2010). Phosphorylated Yki/YAP/TAZ is excluded from the nucleus, preventing it from associating with the transcription factor Scalloped (Sd; mammalian TEAD1-4) to drive expression of growth-promoting transcriptional client genes (VASSILEV *et al*. 2001; WU *et al*. 2008; ZHANG *et al*. 2008), reviewed in (HALDER AND JOHNSON 2011). In summary, the canonical Hippo cascade is Hippo/Mst → Warts/LATS ˧YAP/TAZ+TEAD.

Genetic studies in *Drosophila* and mammalian cells revealed an absolute requirement for Wts/LATS in pathway function, but a partial or context-dependent requirement for Hippo/MST1/2. These observations prompted the discovery of other Ste20-family kinases, but outside the Hippo subfamily, that act redundantly or in parallel to Hippo in specific contexts (MENG *et al*. 2015; ZHENG *et al*. 2015), highlighting the complexity of inputs upstream of Wts/LATS. In most metazoans, the Hippo pathway maintains tissue homeostasis by allowing organs to expand or regress in response to developmental or environmental cues, and its dysregulation contributes to tumorigenesis. Upstream kinases and scaffolds act as tumor suppressors, while YAP/TAZ and TEAD can serve as oncogenic drivers when overactivated (MOROISHI *et al*. 2015; PEARSON *et al*. 2021; HAN *et al*. 2024; HARVEY AND TANG 2025).

A distinctive feature of the Hippo pathway that distinguishes it from other pathways is the absence of a conventional extracellular signaling ligand and transmembrane receptor. Instead, the upstream Hippo kinase is activated by diverse structural elements of the cell: the cytoskeleton, apicobasal cell polarity, and substrate-cell/cell-cell adhesion (MARTIN-BELMONTE AND PEREZ-MORENO 2011). Attenuation of these structures accompany epithelial-to-mesenchymal transition, and hence activation of YAP/TAZ signaling is a major driver of tumor progression (KAPOOR *et al*. 2014; LIN *et al*. 2023). Triggering of downstream YAP activity also suppresses anoikis (FRISCH *et al*. 2013).

Yet in the nematode worm *Caenorhabditis elegans*, development is largely invariant and mosaic: most cell fates are determined by lineage rather than by cell-cell signaling. This rigid developmental program underlies its consistent organ size and stereotyped embryogenesis, unlike the plastic growth observed in flies or vertebrates (HARIHARAN 2015; ROTHMAN AND JARRIAULT 2019). Consequently, canonical Hippo phenotypes – tissue overgrowth or undergrowth – have not been observed in *C. elegans*, and conventional forward genetic screens have not recovered mutants in Hippo, Warts, YAP, or TEAD. Nonetheless, molecular conservation suggested that a core Warts→YAP/TEAD module might be present. Warts is conserved as WTS-1, whose loss causes early larval arrest with compromised intestinal integrity under nutrient stress (CAI *et al*. 2009; KANG *et al*. 2009). YAP-1 and the TEAD ortholog EGL-44 physically interact and are negatively regulated by WTS-1 (IWASA *et al*. 2013), and WTS-1-dependent arrest requires both YAP-1 and EGL-44 (LEE *et al*. 2019). Similarly, NFM-1, the ortholog of Merlin/NF2, the well-established co-regulator of Wts in *Drosophila* and a tumor suppressor in mammals, acts in parallel to WTS-1 to support intestinal polarity and developmental progression (LEE *et al*. 2019). Together, these findings suggest that while the molecular framework of WTS-1→YAP-1/EGL-44 is conserved, it does not drive proliferative phenotypes as in other metazoans.

In *C. elegans,* the role of the namesake Hippo kinase in *C. elegans* remains unclear. Two nearly identical Ste20-family kinases, CST-1 and CST-2, share homology with Hippo but differ at the C-terminus: CST-2 lacks three exons encoding the canonical SARAH coiled-coil domain, a motif characteristic of Hippo, Salvador, and RASSF proteins that mediates homo- and -hetero-dimerization and pathway activation (KARCHUGINA *et al*. 2021). RNAi depletion of both genes induces autophagy and affects aging (LEHTINEN *et al*. 2006; WILKINSON *et al*. 2015), and double deletion causes mild neural defects but no arrest, with no clear connection to the WTS-1→YAP-1/EGL-44 axis. These findings have led to the proposal that *C. elegans* lacks Hippo input to WTS-1 (YANG AND HATA 2013).

This background raises a central question: does *C. elegans* employ Hippo-related kinases to activate WTS-1 repression of YAP-1, or has regulation of the pathway diverged from canonical mechanisms? To address this, we developed a fluorescent YAP-1 reporter to directly monitor pathway activity and assess the contributions of CST-1/2 and other Ste20 kinases to WTS-1 regulation in two distinct tissues. We validated this tool by showing that WTS-1 cell autonomously represses YAP-1 in both epithelium and intestine. In the epithelium, Hippo-like proteins CST-1/2, like WTS-1, repress YAP-1, indicating likely conservation of the canonical Hippo signaling pathway. In contrast, in the intestine repression of YAP-1 upstream of WTS-1 may occur independently of or in parallel to CST-1/2. These findings highlight both the conservation and divergence of Hippo signaling mechanisms in *C. elegans*.

## RESULTS

### WTS-1 is required for larval development and represses YAP-1 and EGL-44/TEAD to promote developmental progression

A prior study reported that the *wts-1(ok753)* deletion allele causes early larval arrest (KANG *et al*. 2009). To more comprehensively characterize the terminal phenotype and expand available genetic tools, we analyzed two additional alleles: *wts-1(tm4081)*, a deletion not available during the original study, and *wts-1(re436)*, a gene disruption (see below). We also developed a conditional depletion strategy using an auxin-inducible degron (AID*) to acutely remove endogenous WTS-1 protein (**Fig. S1A**).

As expected, animals homozygous for *wts-1(ok753)* or *wts-1(tm4081)* exhibited early larval arrest. Both alleles were outcrossed and rebalanced with a fluorescently tagged *tmC18* balancer (DEJIMA *et al*. 2018) covering the *wts-1* locus on Chromosome I. Homozygotes segregated from heterozygous mothers arrested at the L2 stage (**Fig. 1A,B**; **S1A**). Arrested animals continued to move and pump for several days, indicating they were alive (**Fig. 1C**). Differential interference contrast (DIC) microscopy revealed no overt structural defects in L2-arrested animals of either genotype. However, both locomotion and pumping declined by Day 3, suggesting reduced physiological health. This finding is consistent with earlier observations of loss of intestinal integrity and cytoskeletal disorganization in *wts-1(ok753)* animals subjected to prolonged starvation (KANG *et al*. 2009). We speculate that sustained WTS-1 loss compromises animal resilience, perhaps including intestinal integrity. This differs from arrest caused by loss of the small GTPase RHEB-1 or mTORC1, where animals remain viable and can live a full *C. elegans* lifespan after hatching (DUONG *et al*. 2020).

**Figure 1.**
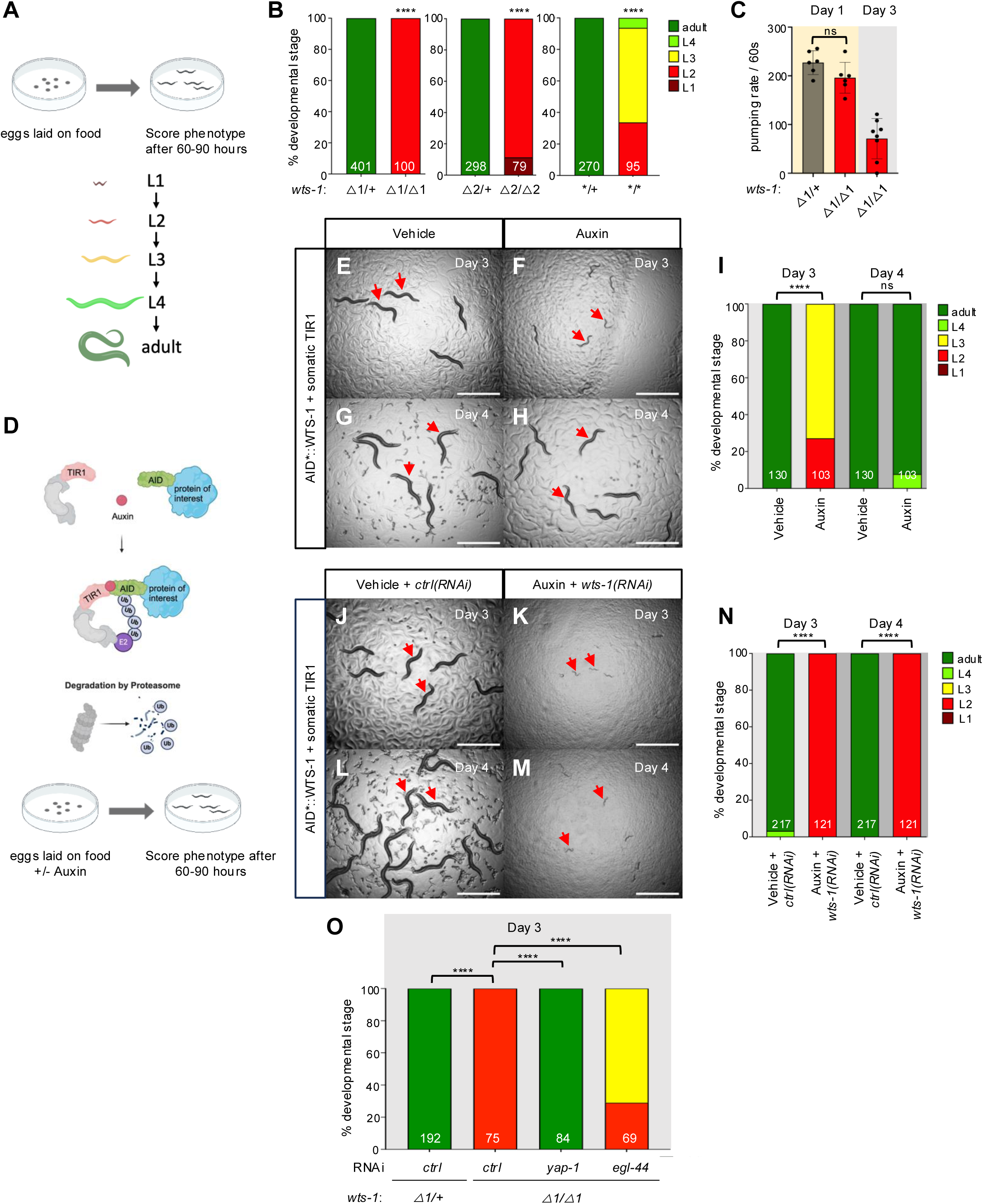
Loss of WTS-1 confers YAP-1/YAP and EGL-44/TEAD-dependent developmental arrest. **(A)** Schematic of experimental design for scoring *wts-1* mutant developmental stages. **(B)** Quantification of arrest stages of homozygous *wts-1* mutant compared to the heterozygote on Day 3. △1 = *wts-1(tm4081).* △2 = *wts-1(ok753).* * = *wts-1(re436[STOP-IN*]). **(C)** Pumping rate/60s of *wts-1(tm4081)* homozygote vs. heterozygote. (**D)** Schematic of protocol for auxin treatment of **E-H)** *wts-1(re419[mTurq2::2xMyc::AID*::wts-1])*; *ieSi57[eft-3p::TIR1::mRuby]* animals grown on vehicle or 1 mM auxin, scored on Day 3 or Day 4. **I)** Quantification of developmental stage for animals in **E-H**. **(J-M)** The same animals as above grown on vehicle or 1 mM auxin and *ctrl(RNAi)* vs *wts-1(RNAi)*, scored on Day 3 or Day 4. *ctrl(RNAi) =* luciferase sequences with no homology in the *C. elegans* genome (Shin *et al*. 2018). **(N)** Quantification of developmental stage for animals in **J-M**. **(O)** Quantification of developmental stages on Day 3 of *wts-1(tm4081)* treated with *ctrl(RNAi)*, *yap-1(RNAi)* or *egl-44(RNAi)*. ****=P<0.0001, ***=P<0.001, **=P<0.01, ns = not significant (Fisher’s exact test).

Knockout consortium mutants are heavily mutagenized and can harbor background mutations. SO using CRISPR/Cas9-dependent genome editing, we introduced a STOP-IN cassette (WANG *et al*. 2018) into the 5’ end of the *wts-1* locus to generate *wts-1(re436)*, a 43 bp insertion that also removes the A of the ATG start codon (**Fig. S1A**). While *wts-1(re436)* mutants arrested, many progressed beyond L2, suggesting that this allele is not a complete loss-of-function (**Fig. 1B**). To assess residual gene activity, we compared *wts-1(re436)* and *wts-1(ok753)* animals grown on *control(RNAi)* versus *wts-1(RNAi)*. *wts-1(ok753)* animals showed mildly enhanced arrest on *wts-1(RNAi)*, with some animals arresting in L1 (**Fig. S1B**), likely due to depletion of maternally contributed gene product. These results suggest that *wts-1(ok753)* is a strong loss-of-function allele, and that a full null phenotype may manifest as L1 or L1/L2 arrest. *wts-1(re436)* animals exhibited a more severe phenotype when exposed to *wts-1(RNAi)* than to *control(RNAi)*, confirming that *re436* mutants retain partial function. As *wts-1* has an 81 nt 5′ UTR, transcriptional initiation is likely unaffected by *re346*. We speculate that the mutant transcript may undergo cryptic translation initiation downstream of the disrupted ATG.

To enable conditional chemical-genetic depletion of endogenous WTS-1, we used CRISPR/Cas9 to insert an *mTurquoise2::2xMyc::AID\** tag at the 5′ end of the *wts-1* coding sequence, using the same guide RNA as the STOP-IN strategy (mTurquois2, or mT2, is a blue fluorescent protein). This minimal AID* degron (44 amino acids) is recognized by the TIR1 E3 ubiquitin ligase, which mediates degradation of AID*-tagged proteins in the presence of auxin (indole-3-acetic acid; IAA; (ZHANG *et al*. 2015). The insertion was determined to be error-free by Sanger sequencing. However, we were unable to visualize blue-tagged WTS-1 beyond very faint signal from the intestinal brush border and we have struggled to detect Myc tags in our lab (Wu and Reiner, unpublished results).

AID*::WTS-1+somatic TIR1 embryos hatched on auxin plates showed delayed development relative to vehicle-treated controls at Day 3. However, by Day 4, these animals had still reached adulthood, indicating developmental delay rather than arrest (**Fig. 1D–I**), indicating depletion was incomplete. In contrast, double depletion with both auxin treatment and *wts-1(RNAi)* caused robust L2 arrest at both Day 3 and Day 4, closely resembling the phenotype of strong loss-of-function alleles (**Fig. 1D, J–N**). Expression of TIR1 in the germline did not confer arrest in auxin-treated AID*::WTS-1 animals (**Fig. S1C**), suggesting that WTS-1 is not essential for embryonic development prior to the maternal-to-zygotic transition in gene expression. Additionally, animals expressing somatic TIR1 but lacking the AID* tag developed normally in the presence of auxin (**Fig. S1D–F**), confirming that auxin itself does not impair development.

### WTS-1 is required to repress YAP-1 and EGL-44/TEAD for development progression past beyond L2

Previous work by Lee and colleagues demonstrated that the *wts-1* mutant arrest depends on the activity of YAP-1 and the TEAD transcription factor EGL-44 (LEE *et al*. 2019). This observation is consistent with the conserved role of Warts/LATS kinases in repressing YAP/TAZ and TEAD in *Drosophila* and mammals (*i.e.* Hippo/Mst → Warts/LATS ˧ YAP/TAZ+TEAD) (HANSEN *et al*. 2015; DASGUPTA AND MCCOLLUM 2019). We tested whether RNAi directed against *yap-1* or *egl-44* could suppress the L2 arrest of *wts-1(tm4081)* animals. *yap-1(RNAi)* fully reversed while *egl-44(RNAi)* partially reversed the arrest (**Fig. 1O**). Similarly, both *yap-1(RNAi)* and *egl-44(RNAi)* suppressed the developmental delay caused by auxin treatment in AID*::WTS-1 animals (**Fig. S1G–J**). These results corroborate the findings of Lee *et al*. using new genetic alleles and validate the use of conditional WTS-1 depletion. We conclude that WTS-1 represses YAP-1 and EGL-44/TEAD to permit developmental progression, a mechanism that is conserved among metazoans.

### Generation of tagged endogenous YAP-1 for use as a reporter of upstream signaling

The transcriptional coactivator *Drosophila* Yorkie (Yki) and its human orthologs Yap and TAZ are well established reporters of upstream Wts/LATS activity. Their subcellular localization reflects regulation by upstream kinases, particularly the Warts serine/threonine kinase (LATS1/2 in mammals). In this conserved mechanism, phosphorylation of YAP by Warts prevents nuclear entry by promoting cytoplasmic retention; inactivation of Warts permits dephosphorylation and nuclear translocation of YAP (HANSEN *et al*. 2015; DASGUPTA AND MCCOLLUM 2019).

Hata and colleagues (IWASA *et al*. 2013) identified the *C. elegans* ortholog of YAP-1 and generated a high-copy integrated translational reporter (*yap-1*p>*yap-1::gfp*). GFP expression from this transgene is broad and YAP-1::GFP signal is predominantly cytoplasmic, with pronounced nuclear exclusion, particularly in epithelial tissues. WTS-1 and the 14-3-3 protein FTT-2 are predicted inhibitors of YAP-1 nuclear localization (HANSEN *et al*. 2015; DASGUPTA AND MCCOLLUM 2019). Consistent with this prediction, RNAi-mediated depletion of *wts-1* or *ftt-*2 resulted in nuclear accumulation of YAP-1::GFP, supporting an evolutionarily conserved role for Warts-mediated inhibition of YAP in *C. elegans* (IWASA *et al*. 2013).

In contrast, RNAi of other putative upstream components failed to cause nuclear accumulation of YAP-1::GFP. These negative results include depletion of *cst-1* and *cst-2*, which encode predicted Hippo-like kinases resembling mammalian MST1/2, the eponymous initiators of the pathway. These negative results raise doubts that the Hippo pathway is conserved in *C. elegans* upstream of Warts and Yap (IWASA *et al*. 2013). Indeed, the “Hippo pathway” in *C. elegans* has been debated as to whether it is a conserved signaling module, a collection of disconnected components, or ignored in the discussion altogether (HALDER AND JOHNSON 2011; YANG AND HATA 2013). Similar ambiguity surrounds the *C. elegans* “Hedgehog” pathway, which retains a GLI transcription factor ortholog but lacks most core pathway elements, but in the case of Hedgehog the conclusion is the pathway has been mostly lost during evolution (BURGLIN AND KUWABARA 2006).

We reasoned that the high expression level of transgenic YAP-1::GFP may reduce sensitivity to perturbation of upstream signal. Possible explanations include buffering of phosphorylation due to excessive protein levels, a lower signal-to-noise ratio in detection of fluorescence, insufficient RNAi knockdown to overcome the threshold required for YAP-1 nuclear entry, or potential redundancy of the duplicated *cst-1* and *cst-2* genes (though these duplicated genes are mostly identical at the DNA level, see below).

To generate a more sensitive and physiologically relevant reporter of endogenous YAP-1 dynamics, we used CRISPR/Cas9-dependent genome editing to insert sequences encoding a short linker, mNeonGreen (a photostable, yellow-shifted fluorescent protein; (SHANER *et al*. 2013) and a 2x FLAG epitope at the 3’ end of the *yap-1* coding sequence, generating *yap-1(re269[yap-1::mNG::2xflag])*; **Fig. S2A**). The resulting edited *yap-1* was error-free by Sanger sequencing. Immunoblotting with anti-FLAG antibody confirmed the predicted tagged protein size of 80.6 kDA (**Fig. S2B**). YAP-1::mNG::2xFLAG was crossed with blue-fluorescent histone marker HIS-72::mT2 and imaged, revealing apparently ubiquitous expression relative to the wild type and low background autofluorescence from unedited controls. Representative images of epithelium and intestine are shown (**Fig. S2C**).

### WTS-1 represses YAP-1 nuclear translocation in epithelium and intestine

In all systems studied to date, Warts/LATS1/2 kinases repress YAP nuclear translocation. To test whether this regulatory mechanism is conserved in *C. elegans*, we used our endogenously tagged YAP-1::mNG reporter to monitor subcellular localization in response to WTS-1 depletion.

We constructed a reporter strain expressing YAP-1::mNG (yellow, imaged at 514 nm) and HIS-72::mTurquoise2 (blue, imaged at 445 nm) for nuclear reference. This strain also harbored a *rrf-3* loss-of-function mutation to enhance sensitivity to RNAi (SIMMER *et al*. 2002). Despite this sensitization, *wts-1(RNAi)* did not cause overt developmental delay or lethality in this strain.

However, imaging revealed that *wts-1(RNAi)* robustly induced nuclear accumulation of YAP-1::mNG in both epithelial cells (**Fig. 2A-C**) and intestinal cells (**Fig. 2D-F**), compared to *control(RNAi)*. These results were recapitulated using the AID*::WTS-1 system: addition of auxin induced nuclear translocation of YAP-1::mNG in epithelial (**Fig. S2D-F**) and intestinal cells (**Fig. S2G-I**), relative to vehicle-treated controls.

**Figure 2:**
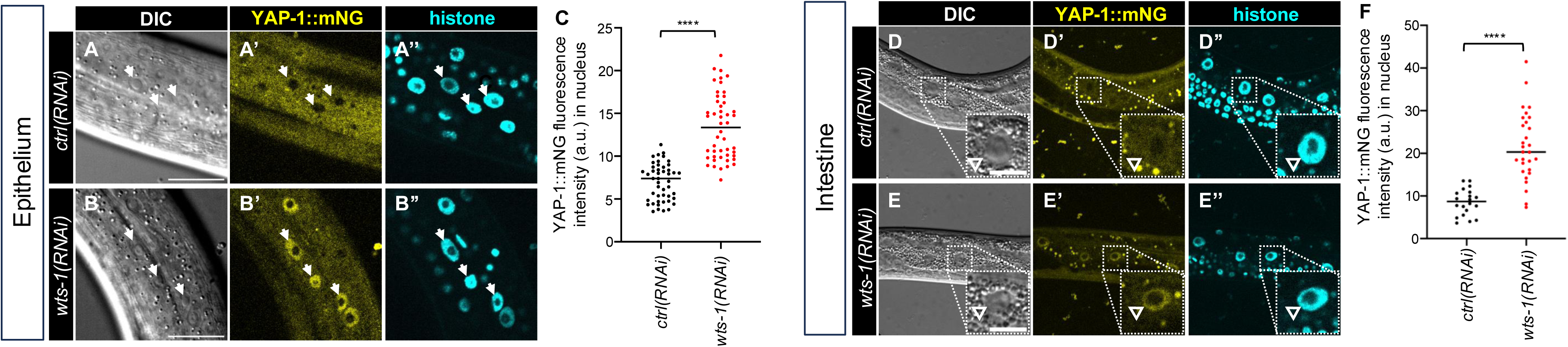
YAP-1::mNG functions as a reporter for WTS-1 activity in epithelium and intestine. **(A-C)** Confocal and DIC photomicrographs and quantification of lateral epithelium; animals are oriented side-up. Animal genotype is *rrf-3(re390[STOP-IN])*; *his-72(erb77[his-72::linker::mTurquoise2])*; *yap-1(re269[yap-1::mNG::2xflag])* to confer hypersensitivity to RNAi. **(A-B’’)** Cytoplasm-to-nuclear translocation of endogenous YAP-1::mNG grown on *ctrl(RNAi)* vs. *wts-1(RNAi)* bacteria, respectively. **(C)** Quantification of fluorescence intensity in **A-B’’** of nuclear YAP-1::mNG in arbitrary units (A.U.). N = multiple nuclei from 9 and 10 animals, respectively. **(D-F)** Confocal and DIC photomicrographs and quantification of intestine, left-right mid-animal plane; animals are oriented side-up. Animals are depicted at lower magnification to show multiple intestinal nuclei. **(D-E”)** Cytoplasm-to-nuclear translocation of endogenous YAP-1::mNG grown on *ctrl(RNAi)* vs. *wts-1(RNAi)* bacteria. Animal genotype is *rrf-3(re390[STOP-IN])*; *his-72(erb77[his-72::linker::mTurquoise2])*; *yap-1(re269[yap-1::mNG::2xflag])* to confer hypersensitivity to RNAi. **(F)** Quantification of fluorescence intensity in **D-E’’** of nuclear YAP-1::mNG as A.U.. N = multiple nuclei from 4 and 5 animals, respectively. Epithelium scale bars = 20 μm, pop out scale bars for intestine = 10 μm. **** = P<0.0001 (*t*-test).

Unexpectedly, we found that the mNG::2xFLAG knock-in into the C-terminus of YAP-1 suppressed the lethality associated with *wts-1* deletion. Double mutants of genotype *wts-1(tm4081)*; *yap-1(re269[yap-1::mNG::2xflag])* developed normally, showing no arrest or delay, and were fully viable and fertile (**Fig. S2J-L**), in stark contrast to *wts-1* single mutants (**Fig. 1**). Nevertheless, YAP-1::mNG strongly accumulated in the nucleus in both epithelial and intestinal cells of *wts-1(tm4081)* mutants (**Fig. S2M-P’**). Lee *et al*. had showed that *wts-1(RNAi)* in an *rrf-3* mutant background conferred arrest, but the endogenous *yap-1* gene in their experiment was unaltered (KANG *et al*. 2009).

Notably, a previous study used an exogenous *yap-1*p>*yap-1::gfp* transgene (IWASA *et al*. 2013), which would not reveal defects in co-transcriptional activity caused by C-terminal tagging. Yet this defect was unexpected, as key functional domains of YAP-1, including the TEAD binding domain (TBD) and WW domains, are located at the N-terminus and central region of the protein, respectively, across diverse species (HILMAN AND GAT 2011; YANG AND HATA 2013; MESROUZE *et al*. 2021).

We conclude that YAP-1::mNG is a reliable reporter of upstream WTS-1 activity, with nuclear localization reflecting loss of WTS-1-mediated inhibition. However, the C-terminally tagged YAP-1::mNG does not appear to function as a transcriptional coactivator; while some YAP/TAZ proteins have a C-terminal PDZ-recognition motif, these are not present in insect YAP proteins nor in YAP-1 (HILMAN AND GAT 2011). Since *yap-1* deletion mutants are superficially wild type (LEE *et al*. 2019; APKEN AND OECKINGHAUS 2021), the transcriptional defect caused by C-terminal tagging of YAP-1 would remain unnoticed unless assayed in a *wts-1* mutant background, where L2 arrest is reversed by loss of YAP-1 function.

### WTS-1 functions cell autonomously to repress YAP-1 in epithelium and intestine

A key test for of cell-cell signaling function is cell autonomy: does a protein function in the tissue manifesting the mutant phenotype or another tissue? Previous results showed that the early larval arrest of *wts-1(ok753)* animals was rescued by transgenic expression of *wts-1(+)* expressed by an intestinal promoter, demonstrating that intestinal expression of WTS-1 is sufficient to support development (KANG *et al*. 2009). Using our conditional chemical-genetic degron, we used tissue-specific expression of the cofactor for AID*-auxin to test the converse question: in which tissue is WTS-1 necessary to support normal developmental progression?

We expressed TIR1, the E3 ubiquitin ligase substrate recognition protein for AID*, specifically in intestinal or epithelial tissues and treated AID*::WTS-1 with auxin vs. vehicle as described (ASHLEY *et al*. 2021). In AID*::WTS-1 animals expressing epithelial TIR1 and grown on *wts-1(RNAi)* to increase robustness, addition of auxin caused mild growth delay relative to vehicle controls (**Fig. 3A-C**). The same outcome occurred without RNAi, though with reduced severity (**Fig. S3A-C**). Degradation efficacy was monitored using degradation of a tissue-specific AID*-tagged nuclear BFP reporter as an internal control. In epithelial TIR1 animals not treated with *wts-1(RNAi)*, auxin induced BFP degradation and nuclear accumulation of YAP-1::mNG in the epithelium (**Fig. 3G-I’**), but not in the intestine (**Fig. 3I-J’**).

**Figure 3.**
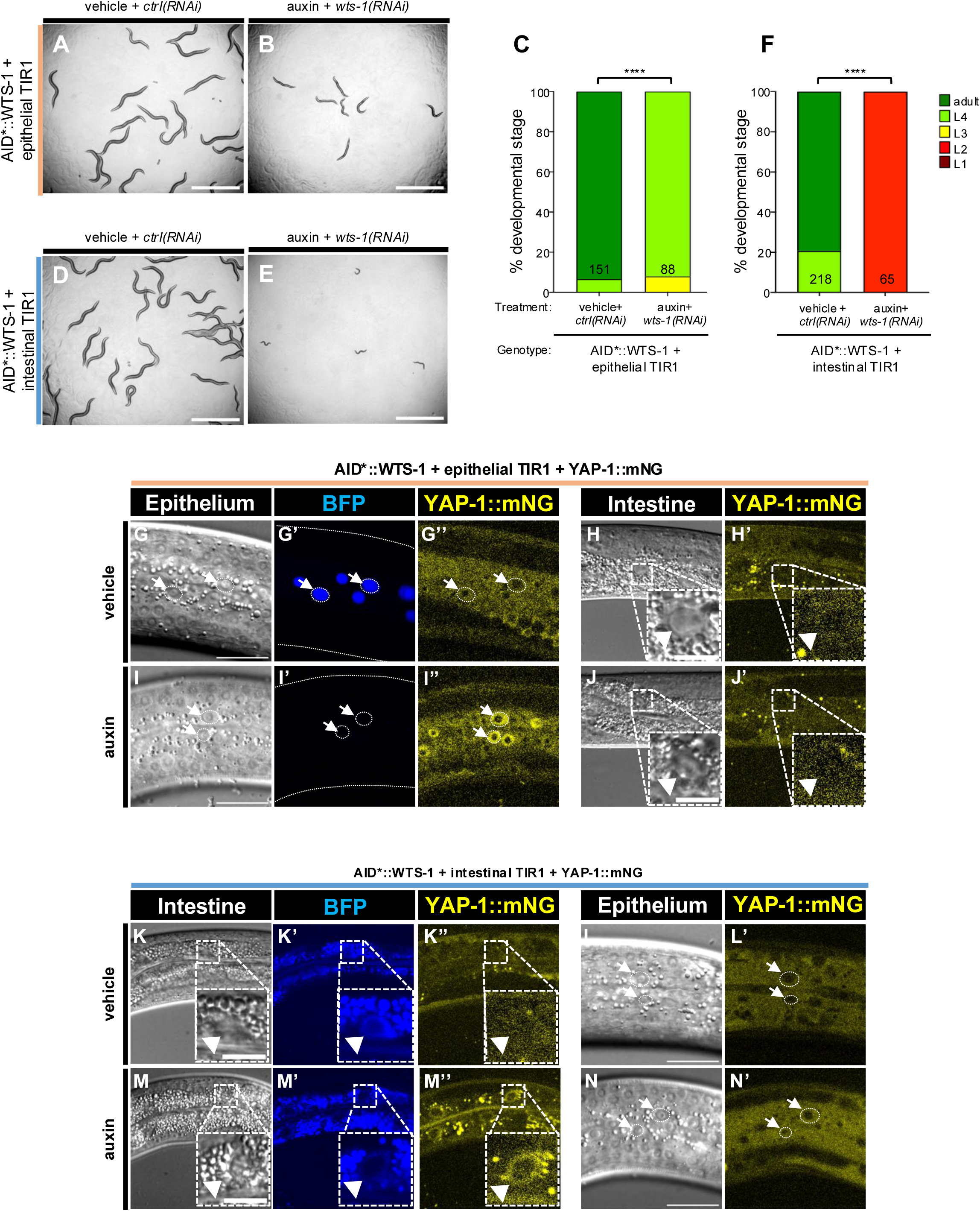
Effects of tissue-specific depletion of AID*::WTS-1. **(A,B)** Bright field photomicrographs of animals with AID*::WTS-1 depletion: global RNAi with epithelial-specific AID* depletion via epithelial-specific expression of TIR1. Genotype is *wts-1(re419[mT2::2xMyc::AID*::wts-1]); reSi2[col-10p>TIR1::F2A::mTagBFP2::AID::NLS::tbb-2 3’UTR]*. **(A)** Animals grown on vehicle+*ctrl(RNAi) vs.* **(B)** auxin+*wts-1(RNAi)*. **(C)** Quantification of **(A** vs. **B)**. **(D,E)** Bright field photomicrographs of animals AID*::WTS-1 depletion: global RNAi with intestinal-specific AID* depletion via intestinal-specific expression of TIR1. Genotype is *wts-1(re419[mT2::2xMyc::AID*::wts-1])*; *reSi12[ges-1p>TIR1::F2A::mTagBFP2::AID::NLS::tbb-2 3’UTR]*. **(D)** Animals grown on vehicle+*ctrl(RNAi) vs.* **(E)** auxin+*wts-1(RNAi)*. **(F)** Quantification of **(D** vs. **E)**. Scale bars = 1 mm. ****=P<0.0001 (Fisher’s exact test). **(G-N’)** Effects on YAP-1::mNG localization of depletion of WTS-1 in epithelium vs. intestine. **(G-J’)** Confocal and DIC photomicrographs of epithelium (**G-G’’,I-I’’**) vs. intestine (**H-H’,J-J’**) with YAP-1::mNG, epithelial-specific TIR1+AID*::BFP and AID*::WTS-1, no RNAi. Genotype is *wts-1(re419[mT2::2xMyc::AID*::wts-1])*; *reSi2[col-10p>TIR1::F2A::mTagBFP2::AI*D::NLS::tbb-2 3’UTR]*; *yap-1(re269[yap-1::mNG::2xFlag])*. **(G-H’)** vehicle-treated animal vs. **(I-J’)** auxin-treated. **(G’** vs. **I’)** Note auxin-dependent depletion of nuclear BFP internal control in epithelium. **(K-N’)** Confocal and DIC photomicrographs of intestine (**K-K’’,M-M’’**) vs epithelium (**L-L’,N-N’**) with YAP-1::mNG, intestinal-specific TIR1+AID*::BFP and AID*::WTS-1, no RNAi. Genotype is *wts-1(re419[mT2::2xMyc::AID*::wts-1])*; *reSi12[ges-1p>TIR1::F2A::mTagBFP2::AID*::NLS::tbb-2 3’UTR]*; *yap-1(re269[yap-1::mNG::2xFlag])*. **(K-L’)** vehicle-treated animal vs. **(M-N’)** auxin-treated. **(K’** vs. **M’)** Note auxin-dependent depletion o nuclear BFP internal control in intestine. Scale bars in epithelial confocal images: 20 μm. Scale bars in intestinal confocal images = 10 μm.

In the converse experiment, intestinal TIR1 expression with auxin treatment in *wts-1(RNAi)* animals caused complete developmental arrest (**Fig. 3D-F**), with weaker effects without RNAi (**Fig. S3D-F**). Auxin treatment also induced intestinal BFP degradation and nuclear translocation of YAP-1::mNG in the intestine, but not in the epithelium (**Fig. 3K-L”, I-J”).**

These results confirm tissue-specific degradation via the AID*-TIR1 system and show that WTS-1 is required in the intestine for developmental progression and repression of YAP-1::mNG. This complements prior rescue studies of *wts-1(Δ)* with intestinally driven *wts-1(+)* (KANG *et al*. 2009). We also find that WTS-1 represses YAP-1::mNG in epithelium, and that epithelial WTS-1 depletion causes a modest growth delay, though whether this is related to intestinal effects remains unclear.

### CST-1/2, the putative *C. elegans* Hippo ortholog, represses YAP-1::mNG in epithelium but not intestine

In *Drosophila* and mammals, Hippo (Hpo) and MST1/2 are Ste20-family kinases (DAN *et al*. 2001; DELPIRE 2009) that regulate growth via activation of Wts and hence repression of YAP/TEAD. While conserved in many species, a clear Hippo ortholog function in *C. elegans* had not been established.

*C. elegans* encodes two Hippo-like kinases, CST-1 and CST-2, are roughly equivalently expressed from tandemly duplicated genes on chromosome X (**Fig. 4A**; mean FPKM of 32 and 24 for *cst-1* and *cst-2*, respectively). These proteins are 100% identical through most of their sequence, including the kinase domain, but differ in the final 13 amino acids of exon 9. *cst-*2 CDS (coding DNA sequence) stops at the end of exon 9, while *cst-1* contains three additional exons that are not present in *cst-2*. Importantly, CST-2 lacks the C-terminal SARAH coiled-coil domain present in all Hpo-related proteins, including CST-1 (**Fig. S4A**), which is essential in other systems for Hippo signaling via homo- and hetero-dimerization (KARCHUGINA *et al*. 2021). Thus, CST-2 may be regulated differently, act independently, or be a non-functional byproduct of genomic duplication.

**Figure 4:**
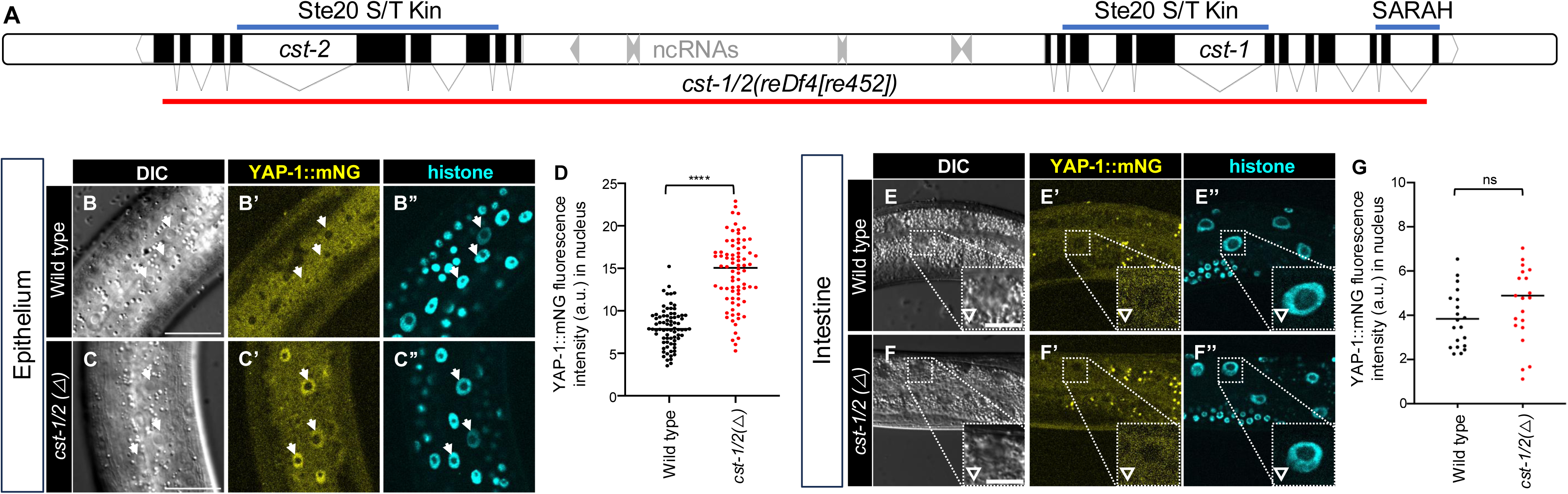
The CST-1/2 putative Hippo orthologs inhibit YAP-1::mNG in epithelium but not intestine. **(A)** Schematic of *cst-1* and *cst-2* genomic region on Chromosome X. The red line indicates the *reDf4[re452]* deletion, which removes *cst-1*, *cst-2* and seven ncRNA genes in between. Lines above the chromosome map indicate conserved protein domains from protein products. CST-2 lacks the C-terminal SARAH (<u>Sa</u>v/<u>RASS</u>F/<u>H</u>po) coiled-coiled dimerization domain present in the C-terminus of *cst-1* and Hippo proteins in other metazoans. **(B-D)** Confocal photomicrographs, DIC photomicrographs and quantification of lateral epithelium (animals are oriented side-up). Animal genotypes are *i.e.* wild type vs. *cst-1/2* deletion: *his-72(erb77[his-72::linker::mT2])*; *yap-1(re269[yap-1::mNG::2xflag]) vs. his-72(erb77[his-72::linker::mT2])*; *cst-1/2(reDf4[re452]) yap-1(re269[yap-1::mNG::2xflag]).* **D)** Quantification of fluorescence intensity in **B-C’’** of nuclear YAP-1::mNG in arbitrary units (A.U.). N = multiple nuclei from 9 and 10 animals, respectively. **(E-G)** Confocal photomicrographs, DIC photomicrographs, and quantification of intestine, left-right mid-animal intestinal plane (animals are oriented side-up). Animals are depicted at lower magnification to show multiple intestinal nuclei. Animal genotypes are *his-72(erb77[his-72::linker::mT2])*; *yap-1(re269[yap-1::mNG::2xflag]) vs. his-72(erb77[his-72::linker::mT2])*; *cst-1/2(reDf4[re452]) yap-1(re269[yap-1::mNG::2xflag])*, *i.e.* wild type vs. *cst-1/2* deletion. N = multiple nuclei from 4 animals each. Epithelium scale bars = 20 μm, pop out scale bars = 10 μm. ****=P<0.0001, ns = not significant (*t*-test)

100% DNA identity extends through all shared exons and upstream sequences of *cst-1* and *cst-2*, even extending several hundred bp upstream of the ATG initiator methionine codon (**Fig. S4B,C**). Examined related Caenorhabditid nematodes encode only a single CST-1 ortholog with a SARAH domain, suggesting a recent duplication event in *C. elegans*.

The *cst-1*-*cst-*2 genomic region was previously deleted as a validation of the transposon-mediated MosDEL deletion technology (FROKJAER-JENSEN *et al*. 2010). In our hands, mutants for the resulting lesion, *basDf1*, are viable, grow slightly slowly, and defective for locomotion. We deleted the *cst-1/2* locus – from exon 1 of *cst-2* to intron 11 of *cst-1* – using CRISPR/Cas9-dependent genome editing, creating the *cst-1/2(reDf4[re452])* allele (**Fig. 4A**). Deletion of *cst-1/2*, like *basDf1*, conferred slow growth and mild locomotion defects, but was not arrested.

The absence of lethality from Δ*cst-1/2* mutants, in contrast to Δ*wts-1*, leads us to speculate that CST-1/2 does not recapitulate the role of WTS-1 throughout the animal. Yet in the *reDf4* mutant, YAP-1::mNG translocated to epithelial nuclei (**Fig. 4B-D**), similar to WTS-1 depletion, suggesting CST-1 and possibly CST-2 activate WTS-1 in epithelium.

Strikingly, *Δcst-1/2* did not cause nuclear YAP-1::mNG translocation in intestine (**Fig. 4E-G**). Given that intestinal *wts-1(+)* rescues *wts-1(ok753)* and that intestinal WTS-1 degradation causes arrest, we propose that CST-1/2 is not necessary for the intestinal WTS-1-dependent event, only the epithelial WTS-1-dependent event. This observation is consistent with studies in flies and mammals, where in certain contexts Hippo/Mst1/2 mutants showed less penetrant phenotypes than Warts/LATS mutants, likely due to kinase redundancy at the level of Ste20 family kinases Hippo/Mst1/2 (MENG *et al*. 2015; ZHENG *et al*. 2015).

### Deletion/depletion of both CST-1/2 and Ste20-family kinase MIG-15 cause WTS-1-like L2 arrest

Previous studies identified Ste20-family kinases from the GCK-I and GCK-IV subfamilies functioning redundantly with Hippo/MST1/2 to activate Wts/LATS1/2 in flies and mammalian cells (MENG *et al*. 2015; ZHENG *et al*. 2015). Specifically, these are the GCK-I and GCK-IV subfamilies of the Ste20 family (DAN *et al*. 2001; DELPIRE 2009). Both consist of N-terminal Ste20 kinase domains and C-terminal CNH (Citron homology) domains, yet they are distinct subfamilies in metazoans. The GCK-I subfamily includes *C. elegans* GCK-2, fly Happyhour, and mammalian MAP4K1,2,3,5. The GCK-IV subfamily includes *C. elegans* MIG-15, fly Misshapen, and mammalian MAP4K4,6,7,8. We previously showed that in *C. elegans* GCK-2 promotes vulval 2° fate and MIG-15 promotes vulval 3° fate (SHIN *et al*. 2018; FAKIEH AND REINER 2025).

We tested whether these Ste-20-family GCK-I and GCK-IV subfamily kinases function redundantly with CST-1/2 to activate WTS-1. In AID*::mNG::2xHA::MIG-15 animals with somatic TIR1, auxin treatment of MIG-15-depleted animals caused mild growth delay (**Fig. 5A-D, I-J**). With same experiment in a *cst-1/2(Δ)* background, auxin-induced MIG-15 depletion caused robust L2 arrest, phenocopying *wts-1* mutants (**Fig. 5E-H, I-J**). Also, as with deficient *wts-1*, MIG-15 and CST-1/2 deficient and arrested animals continued movement and feeding (**Fig. 5K**). This arrest phenotype was reversed by *yap-1(RNAi)* and to a lesser extent by *egl-44(RNAi)* (**Fig. 5L-O**). One interpretation is that MIG-15 also contributes to WTS-1/YAP-1 regulation. Alternatively, the function of MIG-15 could be entirely in parallel to CST-1/2; RNAi of *yap-1* or *egl-44* may suppress effects of mutant *cst-1/2* to retore growth, without impacting the downstream consequences of mutant *mig-15*.

**Figure 5.**
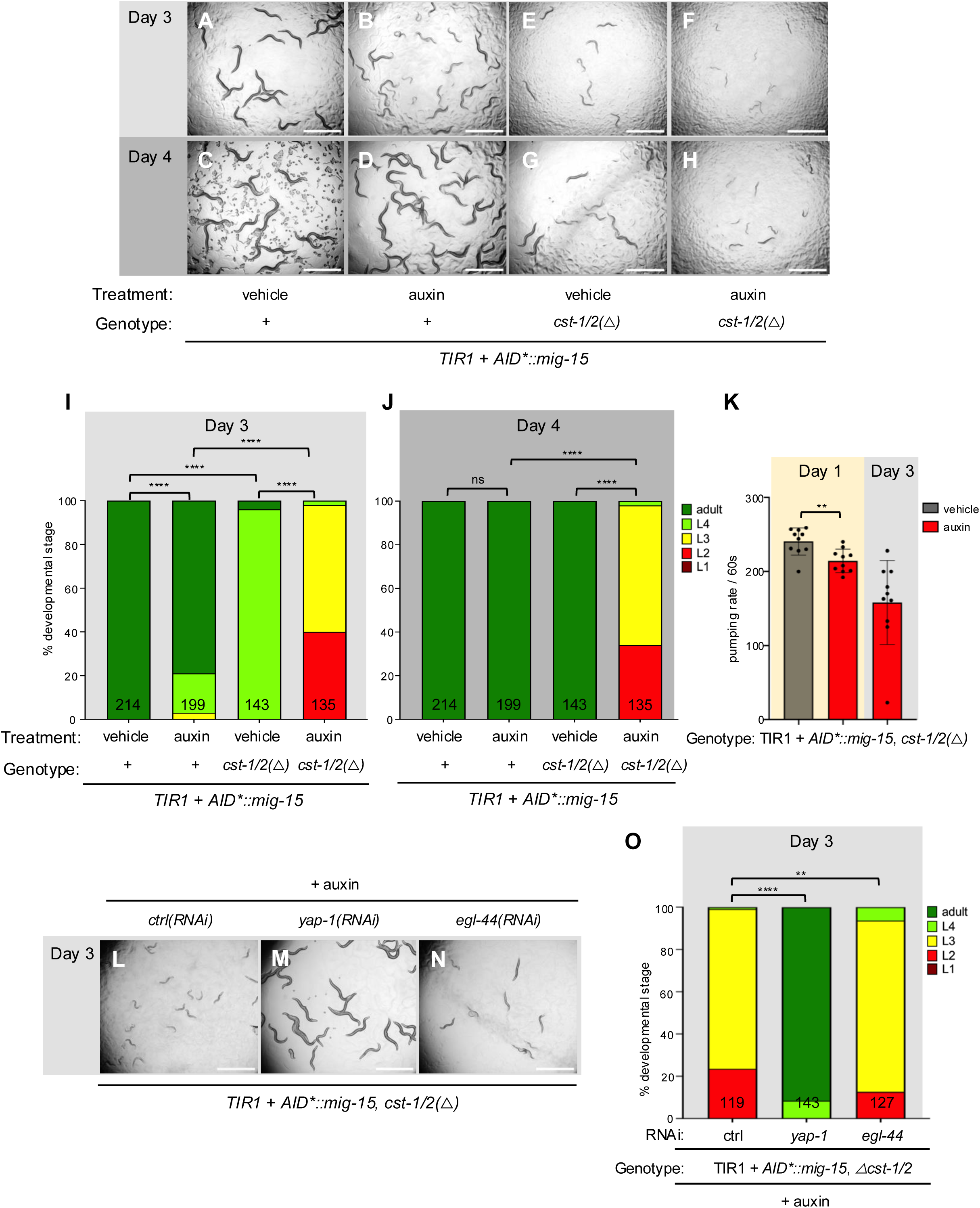
CST-1/2 functions redundantly with MIG-15 to control developmental progression. **(A-H)** Bright-field photomicrographs of depleted AID*::MIG-15 with or without deleted *cst-1/2*, scored on day 3 (**A,B,E,F**) and day 4 (**C,D,G,H**) to illustrate degree of growth delay/arrest. Genotypes are *ieSi57[eft-3p>TIR1::mRuby::unc-54 3’UTR + Cbr-unc-119(+)]; unc-119(ed3); mig-15(re264[AID*::mNG::2xHA::mig-15])* (**A-D**)*vs. ieSi57[eft-3p>TIR1::mRuby::unc-54 3’UTR + Cbr-unc-119(+)]*; *unc-119(ed3)*; *mig-15(re264[AID*::mNG::2x::mig-15]) cst-1/2(reDf1[re484])* (**E-H**). (**B,D** and **F,H**) Treatment with 1 mM auxin *vs.* vehicle on other plates. Depletion of *AID*::MIG-15* causes mild growth delay and locomotion defects (**A/C** vs **B/D**), as published for *mig-15* mutants. Deletion of *cst-1/2* causes mild growth delay and locomotion defects (**A/C** vs **E/G**), as we observed on plates mutant for *basDf1* or *cst-1/2(reDf1[re484])* **(I,J)** Quantification of arrest at Day 3 and Day 4, respectively, reveals the strong arrest of *cst-1/2 mig-15* double mutants relative to single mutants. Animal assays shown in (**E,F,G,H**) and their quantification in (**I,J**), in columns with N of 143 and 135, respectively, are recapitulated in **Fig. S5**, to contrast the effects with and without mutant *gck-2*. All were scored concurrently. **(K)** Pumping rate of control *vs.* double mutant on Day 1 and Day 3 indicates arrest, not lethality. N=10 animals for each group. Animals also continue to swim. **(L-N)** Growth arrest phenotype of AID*::MIG-15 depletion combined with *cst-1/2* deletion was reversed by *yap-1(RNAi)* or *egl-44(RNAi)*. **(O)** quantification of **L-N**. ****=P<0.0001, ***=P<0.001, **=P<0.01, ns= not significant. Scale bar = 1 mm.

To test the Ste20-family GCK-I subfamily member GCK-2, we generated animals with *cst-1/2(*Δ), AID*::MIG-15, and somatically expressed TIR1 with and without *gck-2(re483)*. Animals arrested upon addition of auxin regardless of whether *gck-2* was deleted (**Fig. S5A-J**). Triple mutant *gck-2 cst-1/2 mig-15* animals continued feeding (**Fig. S5K**), like *cst-1/2 mig-15* mutants or *wts-1* mutants, above. A STOP-IN allele, *gck-2(re427)*, edited into the *cst-1/2(*Δ) YAP-1::mNG animal, failed to send YAP-1::mNG into the nucleus (**Fig. 5SL-O**). Thus, GCK-2 does not appear to contribute to this developmental event, though we cannot rule out GCK-2 in other Hippo-regulated signaling events in the animal.

### Ste20-family kinase MIG-15 represses YAP-1::mNG levels but not nuclear translocation

Given the synthetic L2 arrest in auxin+AID*::MIG-15-depleted *cst-1/2(*Δ) animals, we examined YAP-1::mNG localization in these backgrounds. Our MIG-15 tag thus far is with sequences encoding mNeonGreen (FAKIEH AND REINER 2025), which would interfere with visualization of YAP-1::mNG nuclear translocation. Consequently, we used CRISPR/Cas9-dependent genome editing to insert mTurquoise2::2xMyc::AID* at the 5’ end of *mig-15* in animals expressing somatic TIR1 and YAP-1::mNG (**Fig. S6A**). MIG-15 depletion by auxin caused a protruding-vulva phenotype found in all *mig-15* mutants (**Fig. S6B-C**), consistent with strong depletion.

Surprisingly, auxin+AID*::MIG-15 depletion with *cst-1/*2*(+)*increased total YAP-1::mNG levels but did not induce nuclear localization. In epithelial cells, we observed perinuclear aggregation of YAP-1::mNG (**Fig. 6A-C**), potentially indicating localization to Golgi. In intestine, MIG-15 depletion increased YAP-1::mNG expression and weak nuclear localization, but less than that observed in *wts-1* (**Fig. 6D-G**).

**Figure 6.**
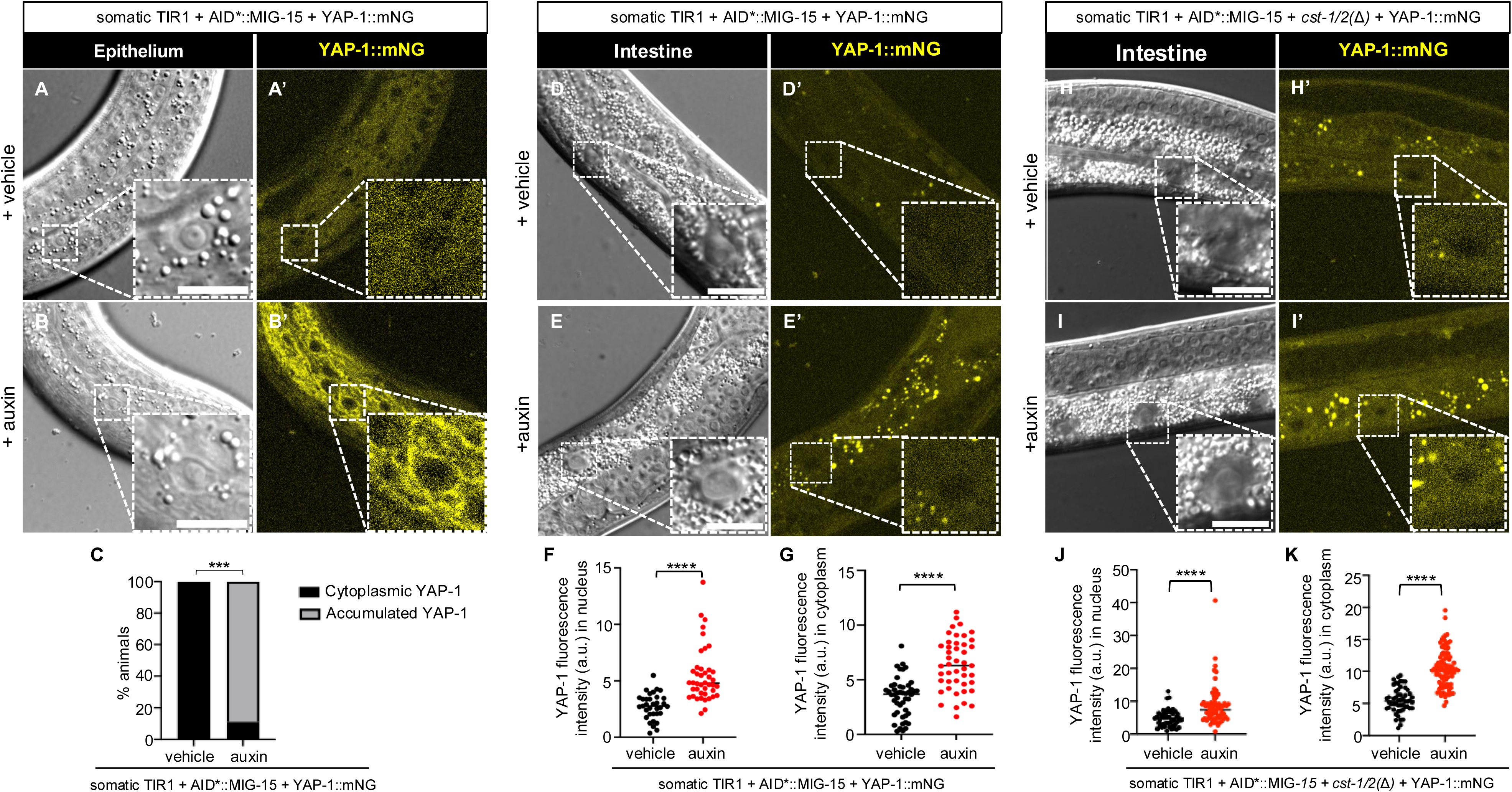
MIG-15 represses YAP-1 levels but does not restrict nuclear translocation. **(A-B’)** Confocal and DIC photomicrographs of YAP-1::mNG in lateral epithelium. Animal genotype is *ieSi57[eft-3p>TIR1::mRuby::unc-54 3’UTR + Cbr-unc-119(+)]; unc-119(ed3)*; *mig-15(re435[mT2::2xMyc::AID*::mig-15) yap-1(re269[yap-1::mNG::2xflag])* on vehicle *vs.* auxin, respectively. **(C)** Quantification of YAP-1::mNG intensity in AID*::MIG-15 animals with and without auxin. N = multiple nuclei from 10 and 9 animals, respectively. **(D-E’)** Confocal and confocal photomicrographs of YAP-1 in midline intestine. Animal genotype is *ieSi57[eft-3p>TIR1::mRuby::unc-54 3’UTR + Cbr-unc-119(+)]; unc-119(ed3)*; *mig-15(re435[mT2::2xMyc::AID*::mig-15) yap-1(re269[yap-1::mNG::2xflag])* on vehicle vs. auxin. Fluorescence intensity was measured for nucleus **(F’)** and cytoplasm **(G’)**, respectively. N = multiple nuclei from 9, 10 animals, respectively.**(H-J’)** Confocal and confocal photomicrographs of YAP-1 in midline intestine. Animal genotype is *ieSi57[eft-3p::TIR1::mRuby::unc-54 3’UTR + Cbr-unc-119(+)]; unc-119(ed3)*; *mig-15(re435[mT2::2xMyc::AID*::mig-15) yap-1(re269[yap-1::mNG::2xflag]) cst-1/2(re479)* on vehicle vs. auxin. Fluorescence intensity was measured for nucleus **(H’)** and cytoplasm **(I**’), respectively. N=10-18 animals. Scale bars = 10 μm. ****=P<0.0001, ***=P<0.001. (*t*-test)

We expected that combined MIG-15 depletion and *cst-1/2(*Δ) would cause robust YAP-1::mNG nuclear localization. Instead, nuclear levels resembled those in MIG-15 depletion alone (**Fig. 6H-K**). While this result was unexpected, the synthetic L2 arrest and rescue by *yap-1(RNAi)* and *egl-44(RNAi)* suggest that MIG-15 and CST-1/2 converge on WTS-1 or YAP-1 regulation. However, they may regulate YAP-1 via different mechanisms or in parallel pathways, as shown in our model of these results (**Fig. 7**).

**Figure 7.**
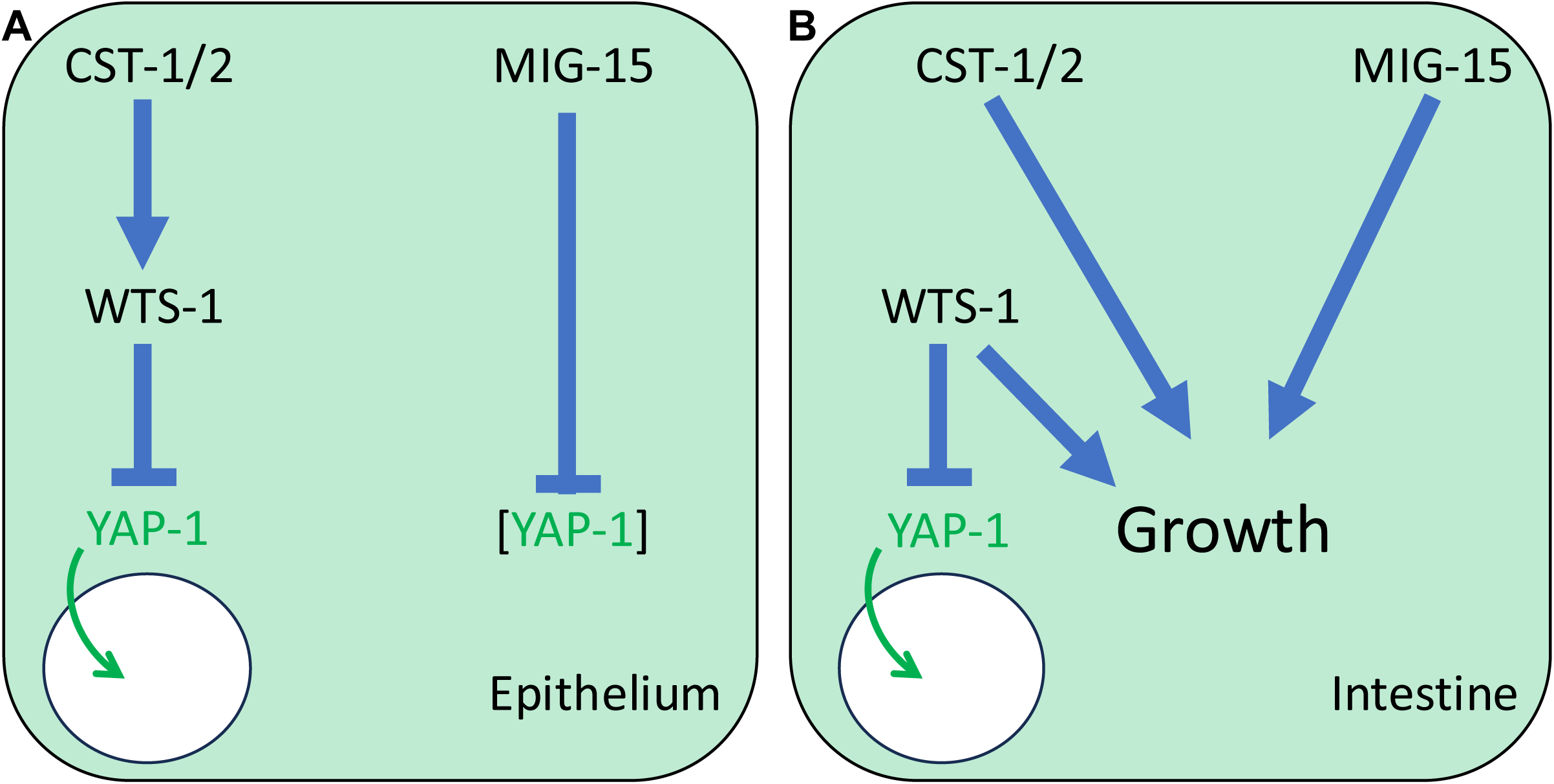
A model for Hippo and Warts signaling in *C. elegans*. **(A)** In epithelium a canonical Hippo pathway functions, with potentially redundant CST-1 and CST-2 phosphorylating and activating WTS-1. WTS-1 in turn phosphorylates and inhibits YAP-1, keeping it out of the nucleus. Disruption of CST-1/2 or WTS-1 results in nuclear translocation of YAP-1, as observed in other systems. Rather than contributing to activation of WTS-1 to repress YAP-1, MIG-15 inhibits YAP-1 protein levels. **(B)** In Intestine a non-canonical Hippo pathway functions. Unknown upstream inputs regulate WTS-1 to phosphorylate and retain YAP-1 out of the nucleus. Loss of WTS-1 causes nuclear translocation of YAP-1 but also L2 arrest, indicated as “Growth.” CST-1/2 and MIG-15 act redundantly to maintain growth.

## DISCUSSION

In this study, we established that *C. elegans* epithelium possesses the molecular wiring of a canonical Hippo→Warts ˧ YAP/TEAD pathway, while also revealing unexpected divergence in its tissue-specific regulation, specific in intestine. Using an endogenous YAP-1 fluorescent reporter, we showed that WTS-1 cell-autonomously excludes YAP-1 from the nucleus in both epithelia and intestine. Tissue-specific depletion of WTS-1 demonstrated that WTS-1 cell-autonomously prevents animal arrest in intestine. At the same time, our data show that the upstream Hippo-related kinases CST-1/2 contribute to YAP-1 regulation only in the epithelium, while intestinal regulation proceeds through a distinct, CST-1/2-independent or CST-1/2-parallel mechanism. We identified MIG-15, the nematode ortholog of *Drosophila* Msn and mammalian MAP4K4,6,7,8 kinases – implicated in activation of fly and mammalian Wts/LATS in parallel to Hippo/MST1/2 – as redundant with CST-1/2 developmental progression past the L2 stage and reliant on inhibition of YAP-1/EGL-44. Unexpectedly, MIG-15 represses of abundance of YAP-1 protein, perhaps through blocking a degradative process, but not nuclear translocation. Together, these results establish a hybrid picture: a canonical Hippo-like cascade in the epithelium *vs.* noncanonical, parallel inputs to WTS-1 in the intestine.

### Divergence of Hippo inputs in *C. elegans*

The presence of one Hippo kinase with a canonical SARAH-domain (CST-1) and another Hippo kinase without (CST-2), coupled with the redundant CST-1/2 kinases not being essential like WTS-1, has fueled uncertainty about whether a *bona fide* Hippo pathway operates in *C. elegans*. Our findings clarify this issue. CST-1/2 act as functional Hippo-like inputs in the epithelium, supporting the model of conservation of the canonical cascade. However, in intestine regulation appears to rely on unknown, alternate kinases. In flies and mammals, MAP4K family members (orthologous to GCK-2 and MIG-15 in worms) act redundantly with Hippo to activate LATS/WTS (MENG *et al*. 2015; ZHENG *et al*. 2015). Our analysis shows that MIG-15, but not GCK-2, alters YAP-1 protein levels in nematodes, suggesting that these kinases may have retained ancillary regulatory functions while losing the ability to control YAP-1 nuclear localization. This divergence highlights the evolutionary plasticity of Hippo inputs across cell types.

Alternatively, MIG-15 and GCK-2 may contribute to phosphorylation of WTS-1 and repression of YAP-1 in other tissues. Epithelium and intestine were assayed because they are large tissues with nuclei readily imaged via confocal microscopy during development. Yet many other tissues or individual cells undergo myriad developmental events while forming the mature animal. For example, EGL-44 regulates identity of a small set of neurons (WU *et al*. 2001), but we would not know if deficiency of WTS-1 or CST-1/2 impacted nuclear translocation of YAP-1 during a key developmental window in the life of these neurons.

### Functional outcomes without proliferation

A striking feature of the *C. elegans* Hippo pathway is the absence of proliferative mutant phenotypes. In other animals, Hippo signaling governs tissue size, regeneration, and tumorigenesis; in nematodes, cell lineages and organ sizes, and indeed animal size, is mostly fixed. Our findings suggest that while the molecular wiring is conserved, the transcriptional outputs of YAP-1/EGL-44 may have shifted. Many YAP/TEAD target genes in *C. elegans* are linked to pathogen responses and gut barrier integrity (MA *et al*. 2020). We hypothesize that YAP-1/EGL-44 has been uncoupled from growth-control genes – either through changes in regulatory DNA elements or altered DNA-binding specificity of EGL-44 – and thus repurposed to support stress and barrier functions. In this sense, *C. elegans* provides a model for Hippo signaling stripped of its growth-control role, emphasizing instead its contributions to tissue robustness.

### Interpreting the arrest phenotype

Loss of WTS-1 causes developmental arrest accompanied by defective intestinal integrity. This arrest could reflect the requirement of an EGL-44/TEAD target gene expression for developmental progression, while other genes are required for tissue integrity. Alternatively, perhaps arrest is a secondary consequence of developmental defects, say perhaps a developmental checkpoint in response to loss of gut integrity, or an intestinal defect that results in starvation. Analogies may be drawn to the mTORC1 pathway, where stress of reduced anabolism can cause programmed diapause at a variety of stages (DUONG *et al*. 2020; SUN *et al*. 2025). Unexpectedly, hyperactivation of YAP-1/EGL-44 upon loss of WTS-1 reduces, rather than enhances, robustness of gut integrity. This finding contrasts with the presumed protective induction of YAP/TEAD in response to pathogenesis/barrier disruption (MA *et al*. 2020), suggesting a feedback loop in which excessive pathway activity destabilizes epithelial function. As with Hippo-mediated growth in other systems, balanced homeostasis may be critical in the gut.

### Tissue- and process-specific roles

Our work complements previous studies showing that RNAi depletion of *cst-1/2* induces autophagy and influences lifespan (LEHTINEN *et al*. 2006; WILKINSON *et al*. 2015), phenotypes not easily explained through WTS-1 or YAP-1. Similarly, Hippo-related roles in neuronal development have been reported in *C. elegans*, unattached to WTS-1 or YAP-1/EGL-44 (FENG *et al*. 2017; JOSEPHSON *et al*. 2017; LEE *et al*. 2018). While our WTS-1 null animals did not exhibit locomotion defects, *cst-1/2* deletions were Uncoordinated and slightly undergrown, suggesting that CST-1/2 may act independently of WTS-1 in sculpting the nervous system. Future experiments will be needed to determine whether YAP-1 or EGL-44 contribute to these locomotory phenotypes.

### Contributions of this work

Our results extend prior studies in several important ways. First, we used a novel endogenous YAP-1 reporter to directly monitor Hippo activity *in vivo*, validating WTS-1-dependent repression across two tissues. Second, we employed tissue-specific depletion strategies to show that WTS-1 regulation is cell-autonomous. Third, we clarified the contributions of CST-1/2, identifying them as functional Hippo-like kinases in the epithelium but perhaps not intestine. Finally, we assessed functions of Ste20-family orthologs, showing that GCK-2 does not contribute to YAP-1 regulation while MIG-15 influences YAP-1 stability but not localization. Together, these analyses position *C. elegans* as a unique model in which Hippo signaling is preserved at the level of WTS-1 inhibition of YAP-1 but diverges in its upstream regulation and biological outputs.

### Broader implications

The field of Hippo signaling has been dominated by studies of over-proliferation, regeneration and tumorigenesis. *C. elegans* provides a contrasting model in which conserved pathway components are retained but uncoupled from growth control. Instead, Hippo signaling contributes to epithelial robustness, stress responses, and possibly neural development. These findings underscore the evolutionary flexibility of Hippo signaling and open new directions for understanding how signaling pathways are rewired to meet the developmental logic of different organisms.

## METHODS

### *C. elegans* handling and genetics

All *C. elegans* strains were derived from the N2 Bristol wild type and grown on NG agar plates seeded with the OP50 strain *E. coli* at 20°C unless otherwise noted. Growth, culturing and nomenclature were generally as described (BRENNER 1974; HORVITZ *et al*. 1979; TULI *et al*. 2018). Wormbase was used extensively during these studies (STERNBERG *et al*. 2024).

### CRISPR/Cas9 mediated genome editing

Guide RNAs were that maximally satisfied a combination of three approaches. First, where possible, G and not T nucleotides were selected at positions −1/−2, or GCGG and not T-TT nucleotides at positions −1/−2/−3/−4 (KAPOOR *et al*. 2014; FARBOUD AND MEYER 2015). Second, we considered strong predicted specificity and efficiency scores using the CRISPOR (http://crispor.tefor.net/) algorithm, which incorporates the original MIT specificity score. Third, we considered strong predicted efficiency scores using the WU-CRISPR (http://crisprdb.org/wu-crispr/) algorithm.

We formulated mixes for microinjection as described previously (FAKIEH *et al*. 2022). Using a dedicated RNase-free bench, we mixed concentrations as final volume per 20 μL: 1 μl of 5 μg/μl stock *Streptococcus pyogenes* Cas9 (PNAbio, #CP01) for 0.25 μg/μl final concentration; 1μl of 2 μg/μl stock universal tracrRNA (IDT) for 0.1 μg/μl final concentration; 1.4 μl each of 0.4 μg/μl stocks of *dpy-10* co-CRISPR and gene-specific crRNAs for 0.028 μg/μl crRNA final concentration each. Not yet at final volume, this mix was incubated at 37°C for 15 min. After incubation, *dpy-10* ssODN repair oligo was added as 3.3 μl of 20 μM stock for final concentration of 3.3 μM; gene-specific column-purified PCR repair template was denatured and renatured as described (GHANTA AND MELLO 2020), and added from stock of ≤300 ng/μl to a final concentration of 100 ng/μl (total final concentration of PCR product no greater than 2 μg/20 μl for reasons of viscosity). As this point, commercial nuclease-free water was added up to 20 μl. (Concentrations before the incubation were calculated per 20 μl, but 20 μl volume was not reached until the final addition of water.)

Adult animals with no more than a single row of embryos in each uterus side were microinjected as described (KADANDALE *et al*. 2009). We used co-CRISPR to generate dominant Rol mutations in F1 animals harboring the *dpy-10(cn64*gf) marker (ARRIBERE *et al*. 2014), which were picked singly or in pools of two to plates. After F1 Rols had laid embryos, the single or pooled parents were picked to PCR tubes, lysed for PCR and detected via triplex PCR. Non-Dpy/Rol F2s were picked singly, incubated overnight to obtain genetic material, lysed and PCR detected to generate homozygous lines. Knockins were sequenced from points in flanking DNA outside that subjected to homology-directed recombination repair. Only expected knockins were used for analysis.

### RNA interference

Plates for bacterially mediated RNAi interference (RNAi) were made with NGM agar supplemented with 1 mM IPTG and 50 μg/mL carbenicillin. Plates were spotted with 80 μl of HT115 bacteria culture expressing dsRNA as described (TIMMONS *et al*. 2001). L4 P0s were picked to 24 hr-seeded RNAi plates, grown overnight, then transferred the next day to fresh RNAi plates for synchronized laying of embryos. The P0s were removed 24 hours later and F1 animals were analyzed at the time appropriate for the experiments. Positive control for efficacy of RNAi plates was *pop-1(RNAi),* where we expect 95-100% embryonic or L1 lethality of the experiment is discarded. For negative control with used *luciferase(RNAi)* as previously described (SHIN *et al*. 2018), to engage the RNAi machinery with a sequence not encoded in the *C. elegans* genome. RNAi was performed at 20°C.

### Pumping assays

Benchtop mechanical clickers were uses to count contractions of the pharynx (“pumping”) per 60 sec. Observations used a Nikon SMZ18 stereo microscope.

### Auxin treatment

Auxin (IAA; Indole-3-acetic acid, 98+%, Thermo Scientific) storage stock solution was prepared at 400 mM in ethanol and stored up to a month at 4°C. For experiments, stock auxin was diluted to 16 mM working auxin stock by diluting storage 400 mM auxin with filtered Milli-Q water. For final concentration of 1 mM auxin in plates, 500 μl of 16 mM auxin was added onto 8 ml NGM agar plates or RNAi plates and allowed to diffuse. For vehicle control, 500 μl of 4% ethanol was similarly added to plates.

### Detection of tagged endogenous YAP-1::mNG::2xFLAG

Mixed-stage animals were washed from 3-4 full but not starved plates using M9 buffer and lysed in 4% SDS loading buffer with boiling at 90°C for 2 min. Samples were run on a 4-15% SDS-PAGE gel (Bio-Rad, #5671084) and transferred to Immobilon-P Membrane, PVDF (EMD Millipore, IPVH00010). Mouse monoclonal anti-Flag antibody (Sigma-Aldrich, F1804) and mouse monoclonal anti-α-tubulin antibody (Sigma-Aldrich, T6199) were diluted 1:2000 in blocking solution (6% w/v non-fat dry milk in PBST). HRP conjugated goat anti-mouse secondary antibody (Sigma-Aldrich, 12-349) was diluted 1:5000 in blocking solution. Chemiluminescent detection was performed using ECL reaction (Thermo Fisher Scientific 32106) and detected via film processor, SRX-101A (Konica Minolta) on X-ray film (Phenix).

### Imaging

Animals used for live imaging were mounted on a 5% agar pad on a glass slide in 5 μl of 2 mg/ml tetramisole in M9 buffer with cover slip. Confocal microscopy used a Nikon Ti2 microscope with a Yokogawa CSU-W1 Spinning Disk, 405 nm, 445 nm, 514 nm lasers, and a Photometrics Prime BSI camera, and NIS Elements Advanced Research software, version 5.42. Quantification of fluorescence intensity were performed by ImageJ (Fiji) with a customized script code (deposited at: https://github.com/ReidTPowell/2025_Genetics_Reiner). In 3-channel confocal photomicrographs analyzed in ImageJ with the script running, the user identifies the center of a nucleus in the DIC channel and placed a point using the “multi-point tool.” The script draws a uniform circle for each point of interest and measures fluorescence intensity. This confers precision in area of circle from sample to sample.

10 cm NG plate photomicrographs were captured using a Nikon Eclipse Ni microscope, 4x objective, ANDOR Zyla sCMOS camera and NIS Elements Advanced Research software, version 4.30, with plate and animals on the stage with sufficient working distance.

### Sequence analysis

Protein sequences were accessed via Wormbase (wormbase.org; (STERNBERG *et al*. 2024) and Alliance of Genome Resources (https://www.alliancegenome.org/). We used the A isoform for both *cst-1* and *cst-2*, thus excluding the two-codon minority splice addition to exon 9 for both. *mig-15* isoform A was selected as the reference and most abundant isoform. Uniprot accession numbers were Q13188 for human MST1/STK3, Q13043 for human MST2/STK4, Q8T0S6 for *Drosophila* Hpo. Protein sequences were aligned with Clustal Omega (https://www.ebi.ac.uk/jdispatcher/msa/).

### Statistical analysis

Animal experiments presented in the same graph were scored concurrently. Statistical analyses were performed by two-tail unpaired Student’s t-test or Fisher’s Exact test (GraphPad Prism 10).

## Software/website

GraphPad Prism was used for statistical analysis and making graphs. BioRender (https://www.biorender.com/) was used to generate illustrations. Exon-Intron Graphic Maker was used for making schematic gene structures http://wormweb.org/exonintron. AI software was used only for text editing purposes.

## Data availability statement

Strains, plasmids and sequence files are available upon request. All data supporting the conclusions of the study and required for reproduction of experiments are present within the text, tables and figures.

## Acknowledgements

We thank L. Vergara in the Center for Advanced Imaging for technical assistance, members of the Reiner lab for helpful discussions and technical expertise, J. Bembenek (Wayne State University) for the strain harboring HIS-72::mT2, M. Bastiani (University of Utah) for *basDf1*, Y. Hata (Tokyo Medical and Dental University) and B. Grant (Rutgers University) for the strain harboring *ihIs35[yap-1*p*>yap-1::GFP], and* S. Mitani (Japanese National Bioresource Project) for *wts-1(tm4081)*. Some strains were provided by the *Caenorhabditis* Genetics Center, which is funded by the NIH Office of Research Infrastructure Programs (P40 OD010440). Wormbase was used extensively. Our CRISPR protocol was based on an amalgam protocol generously shared by Dr. Andy Golden (NIDDK).

## Supplementary Tables

Strains used are shown in Table S1.

Oligonucleotides in Table S2.

Guide RNAs in Table S3.

Plasmids in Table S4.

## Suppe

**Figure S1.**
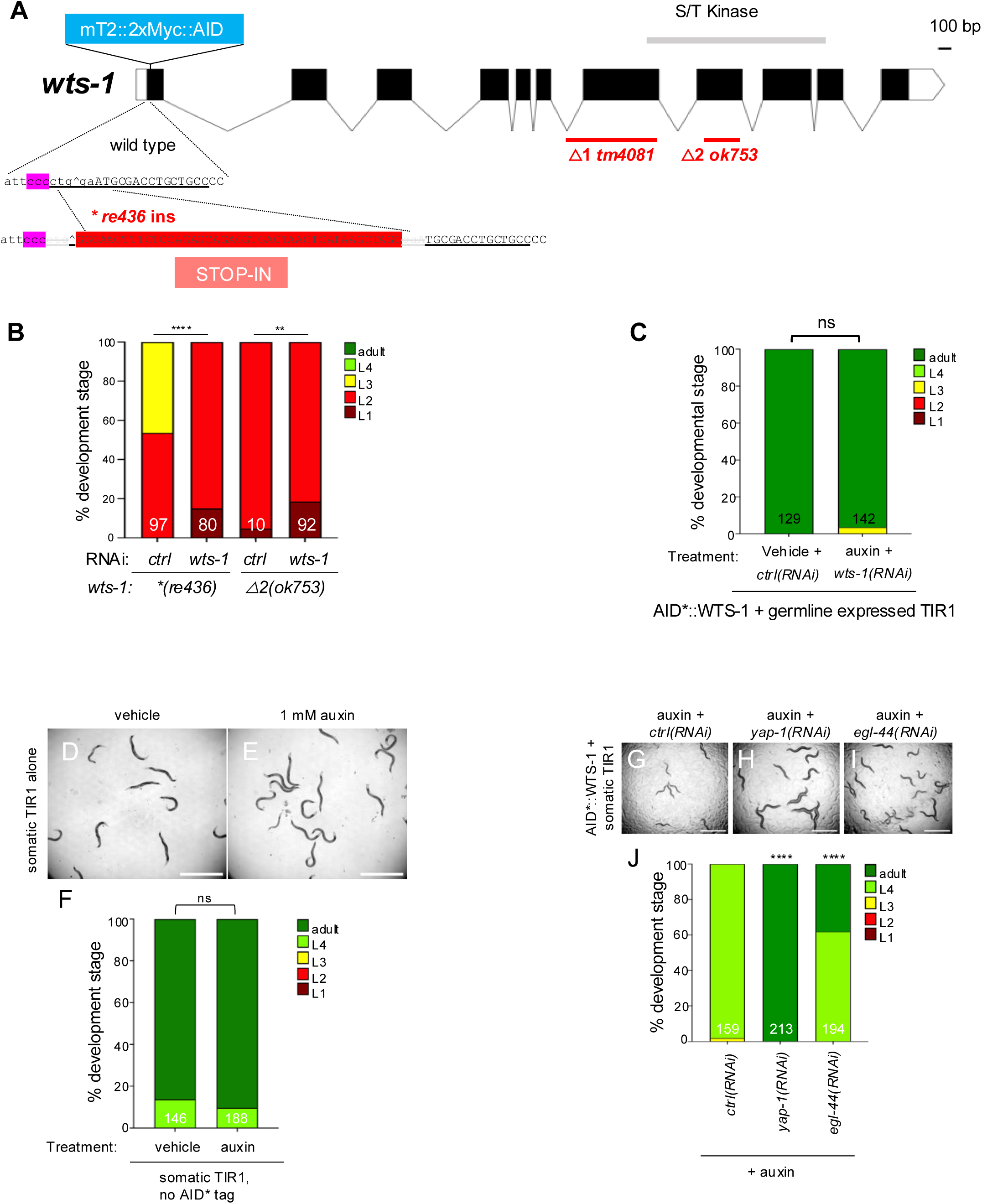

**Figure S2.**
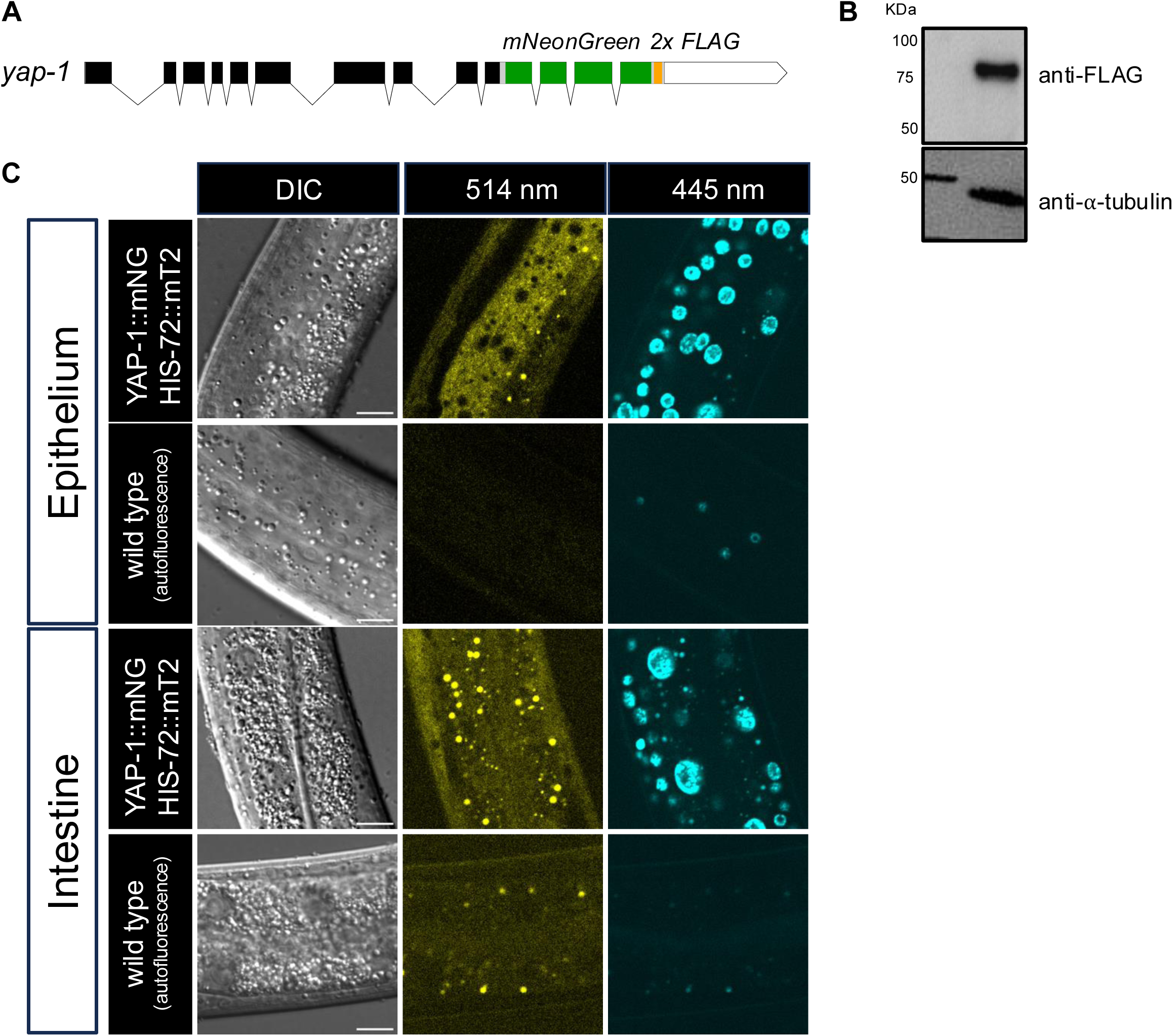
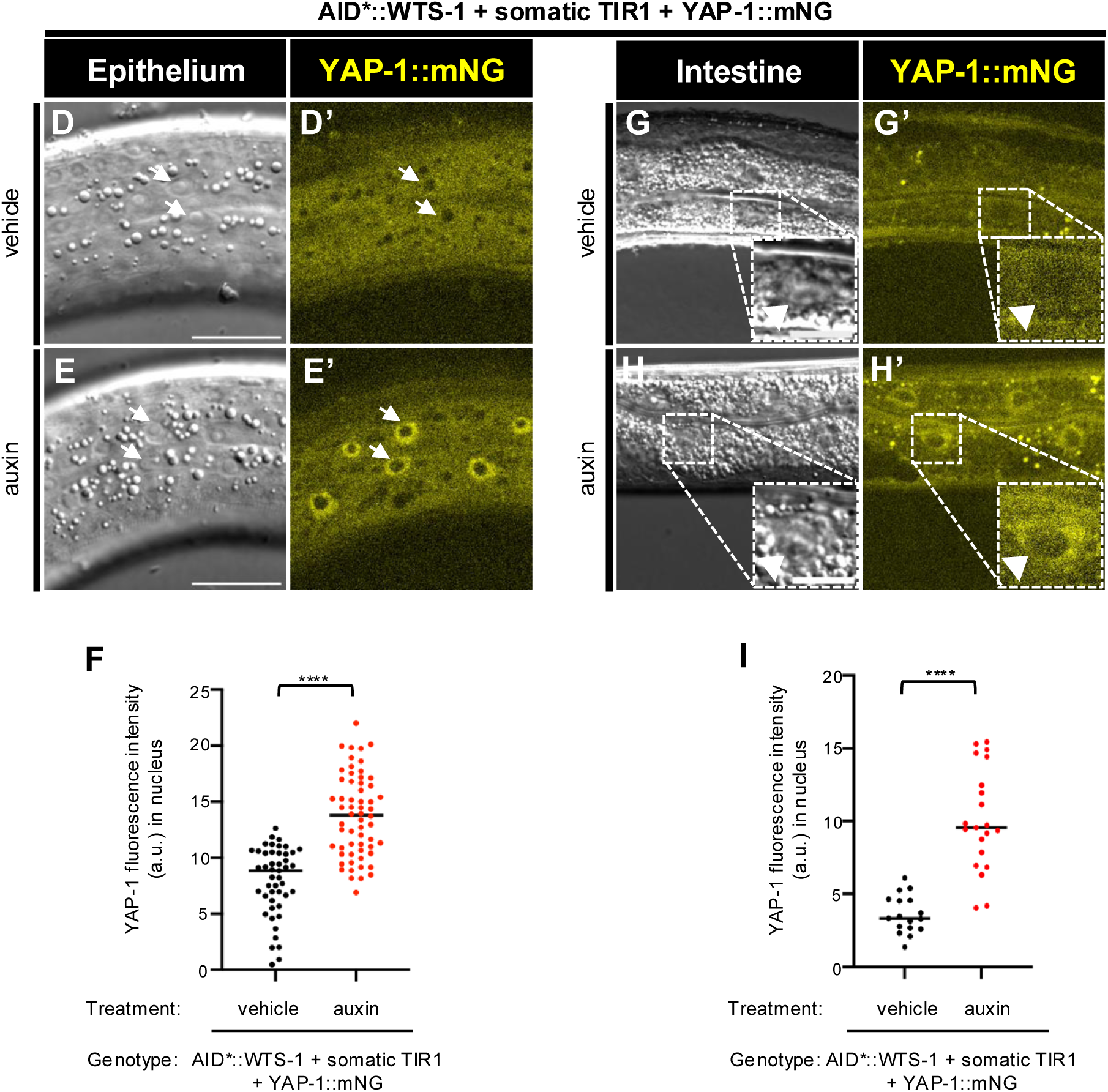
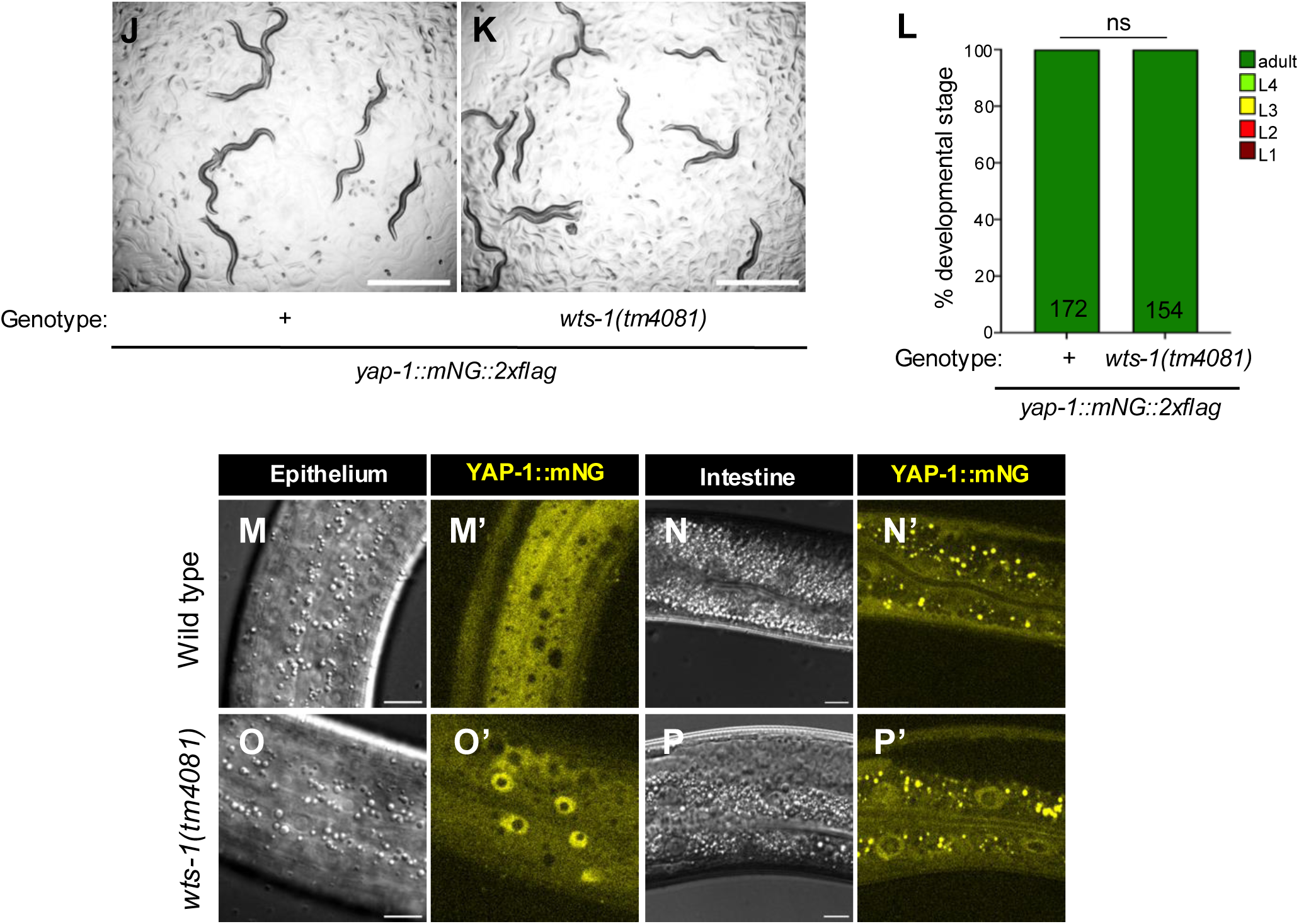

**Figure S3.**
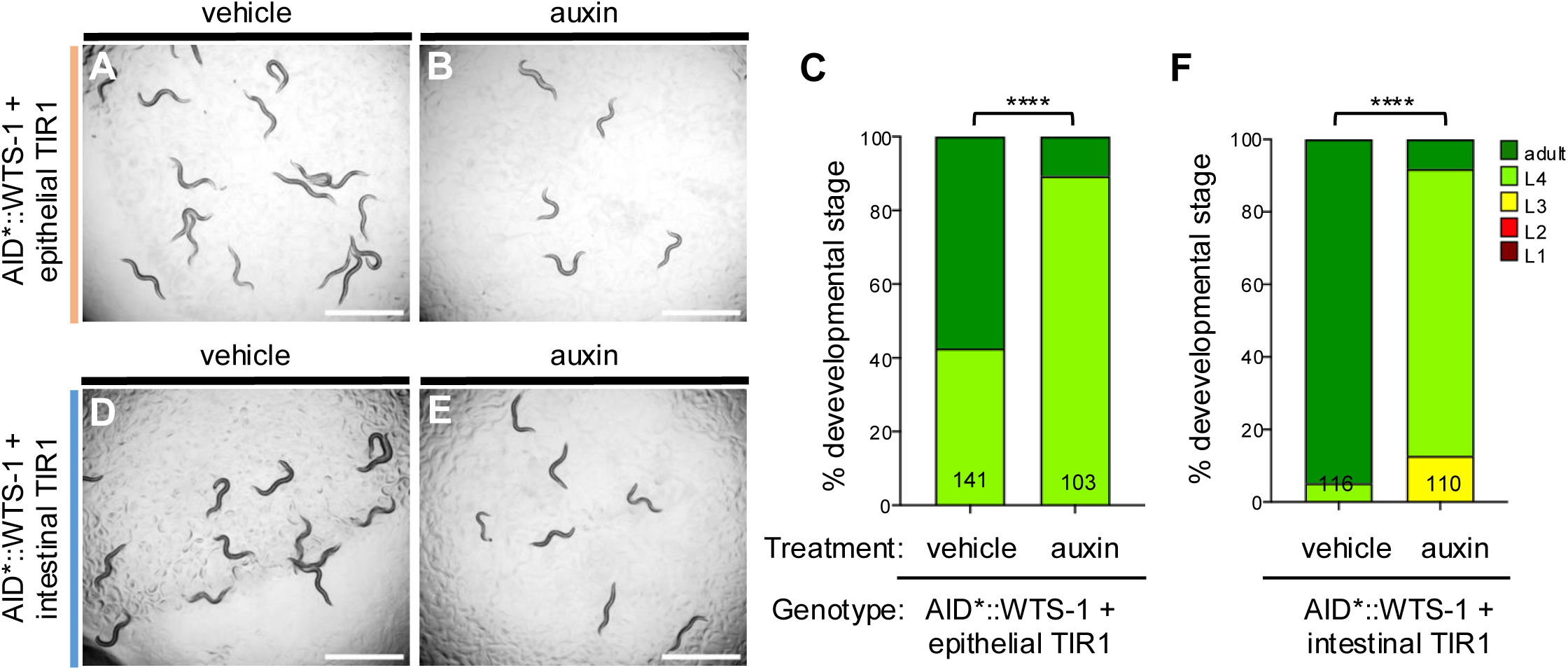

**Figure S4a.**
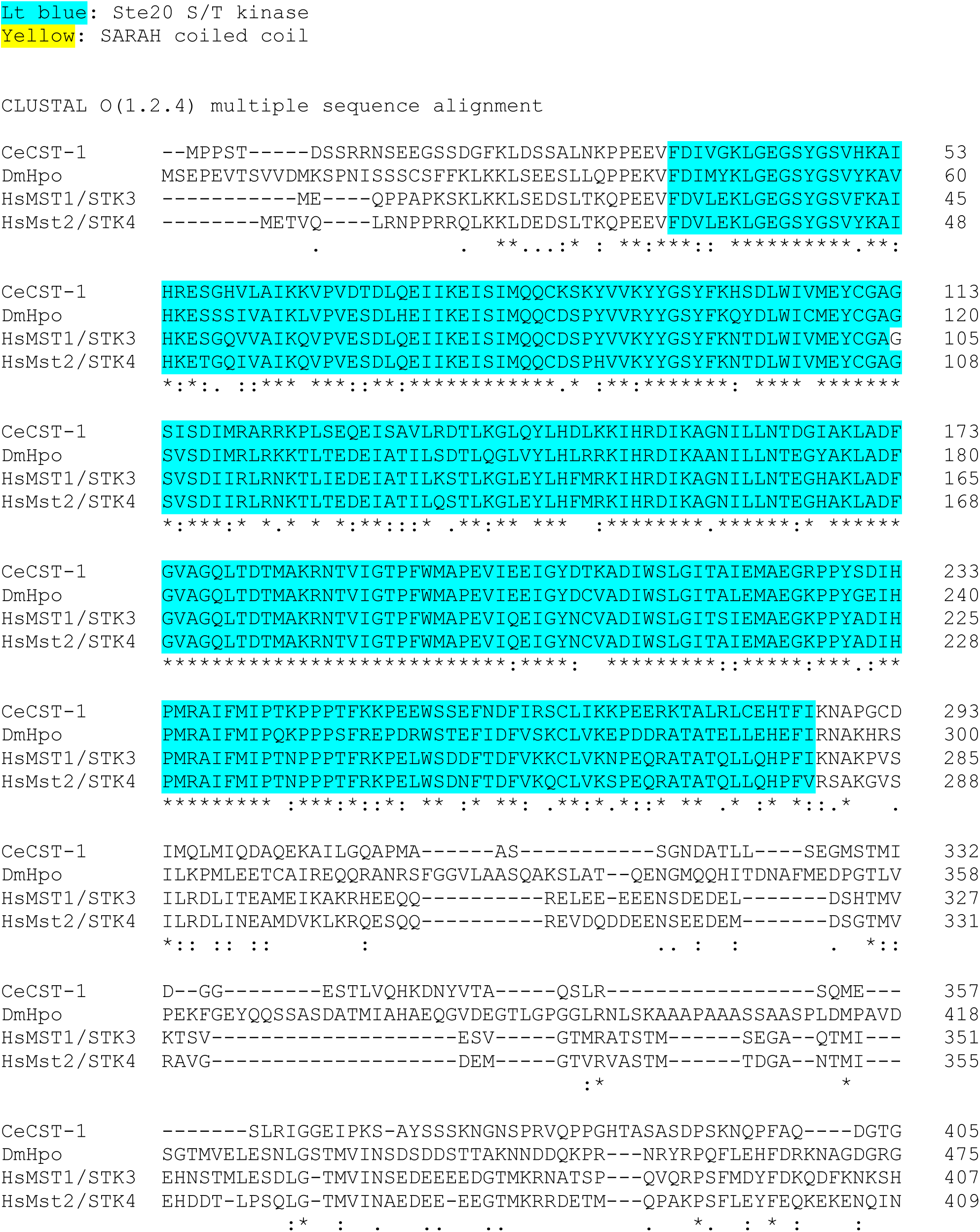
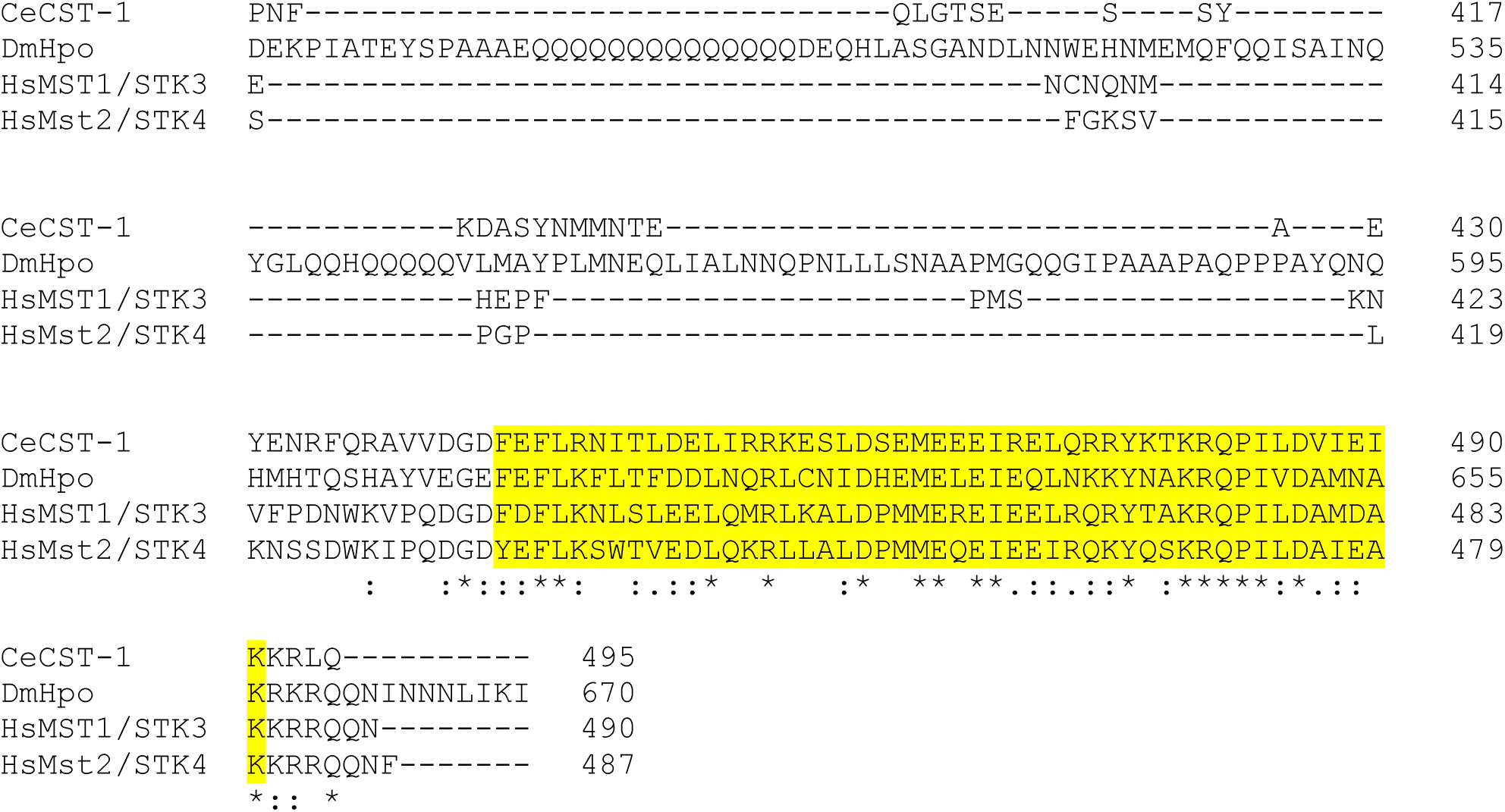

**Figure S4b.**
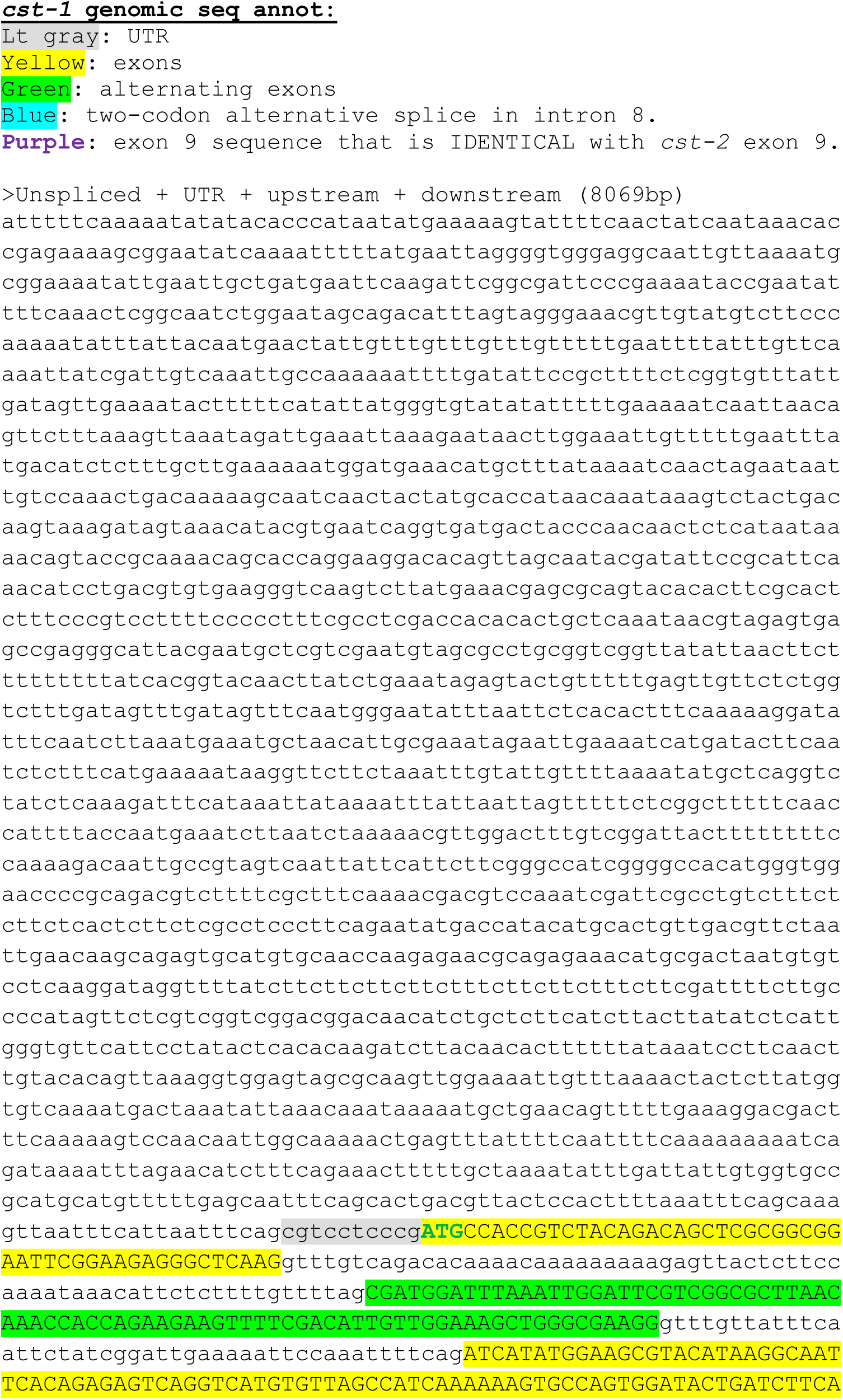
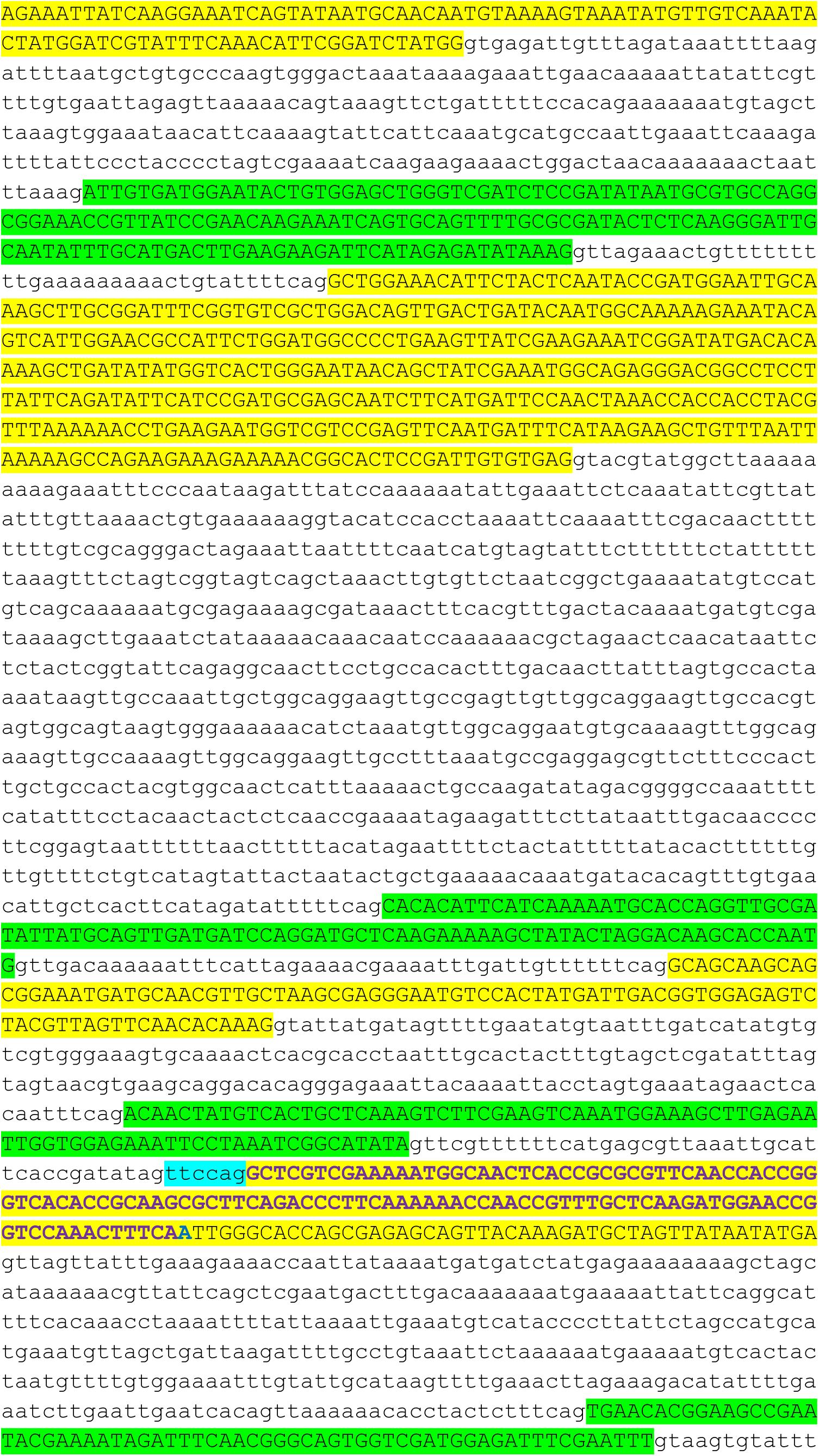
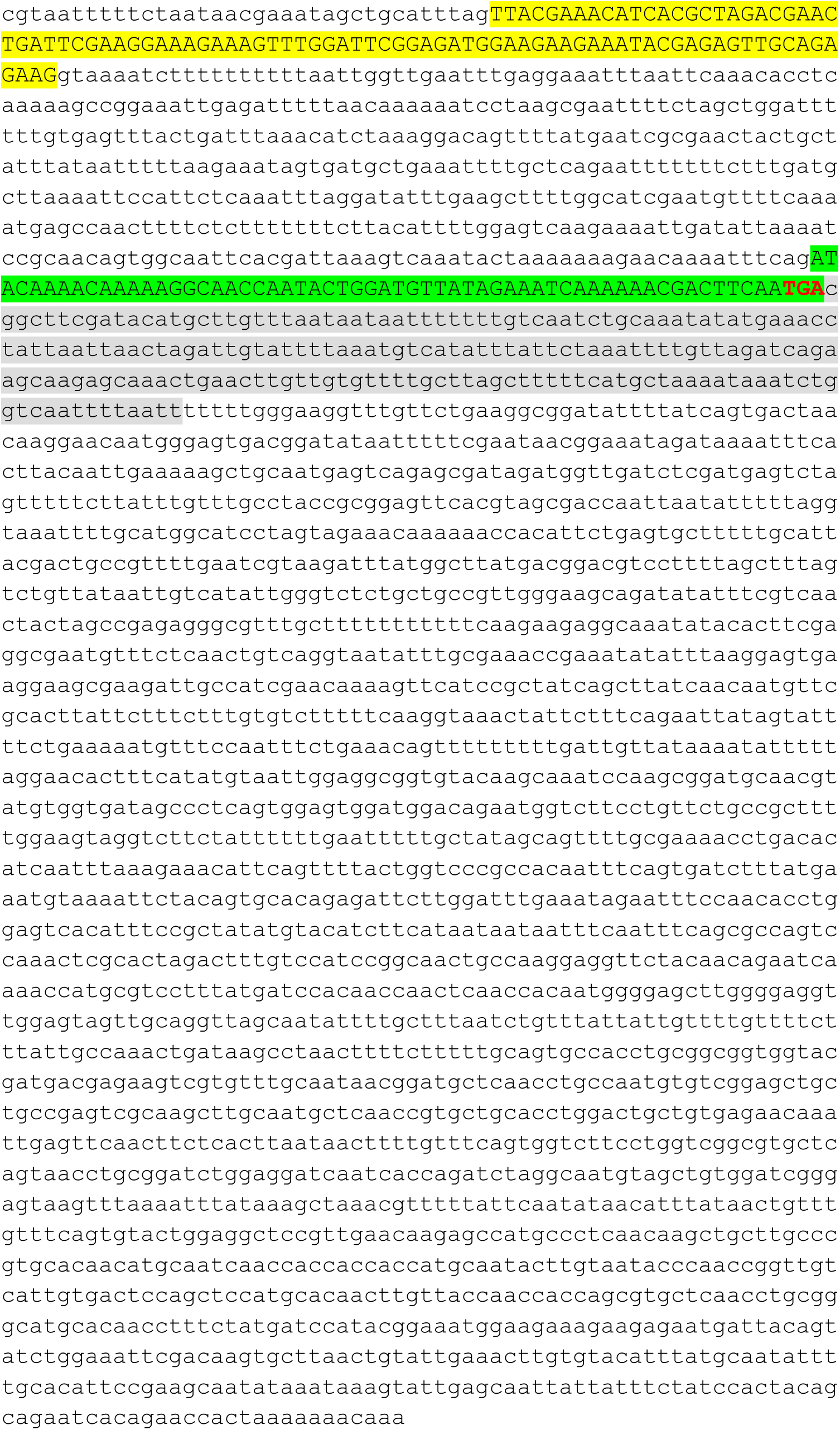
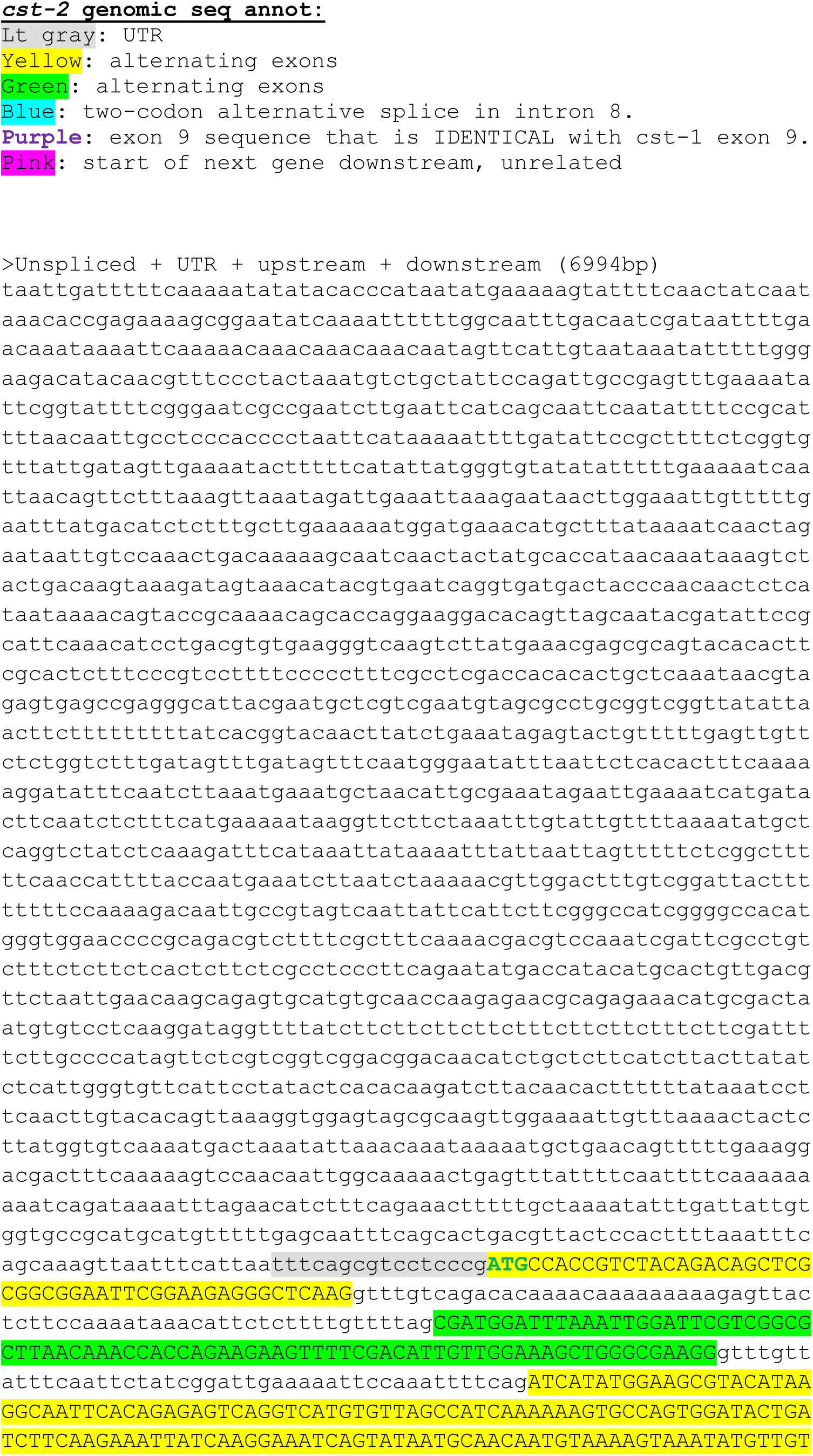
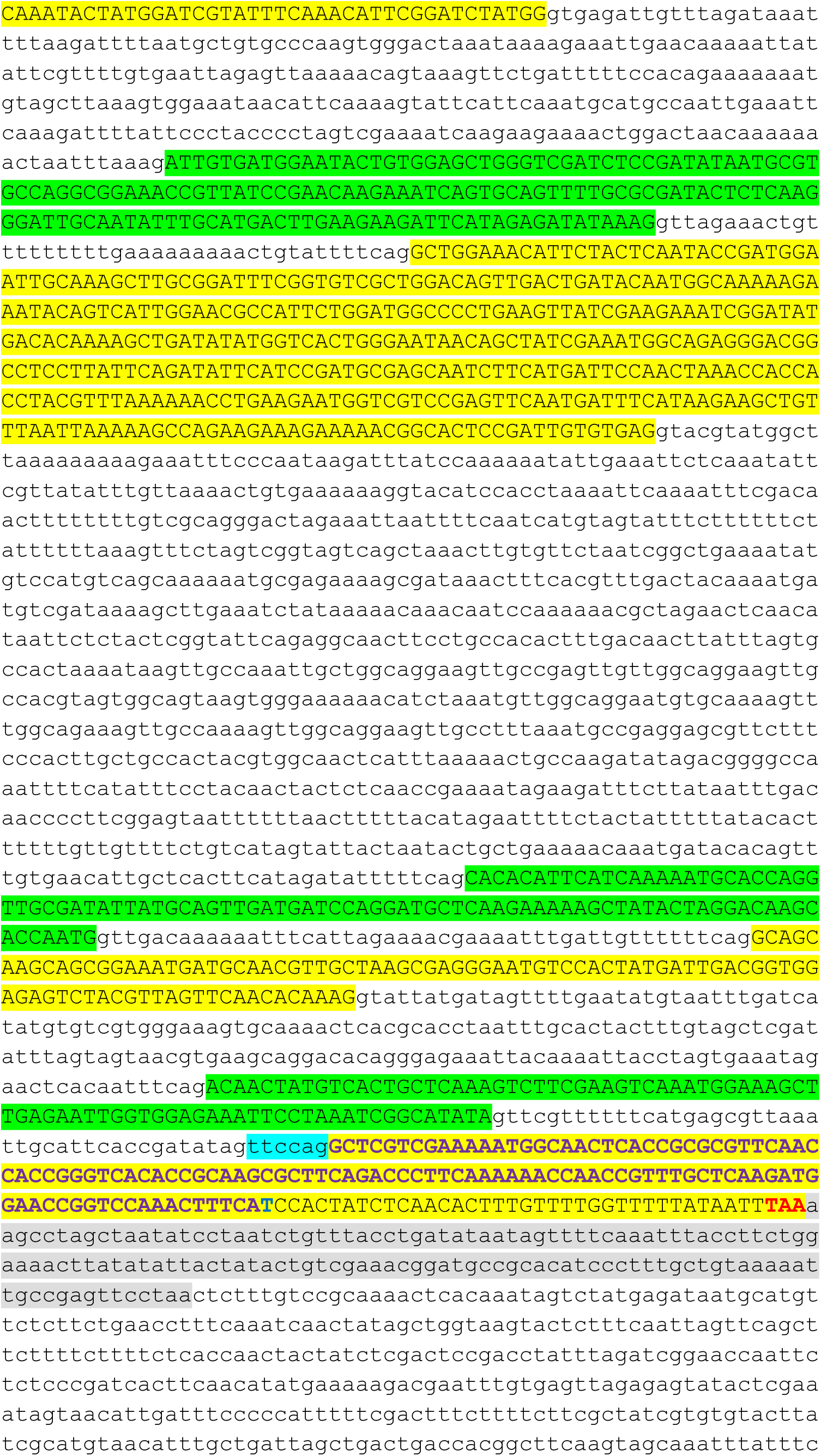
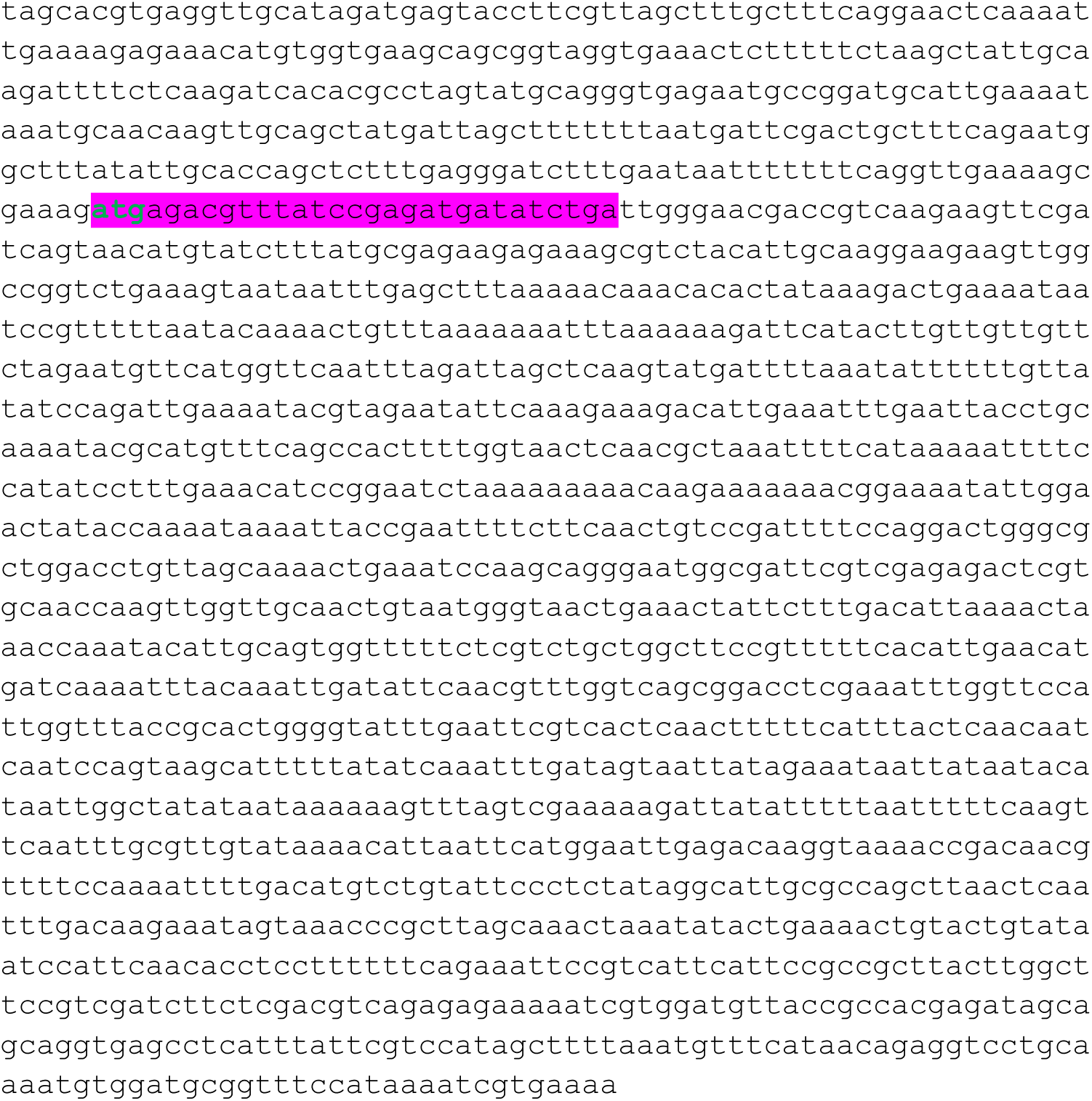

**Figure S5.**
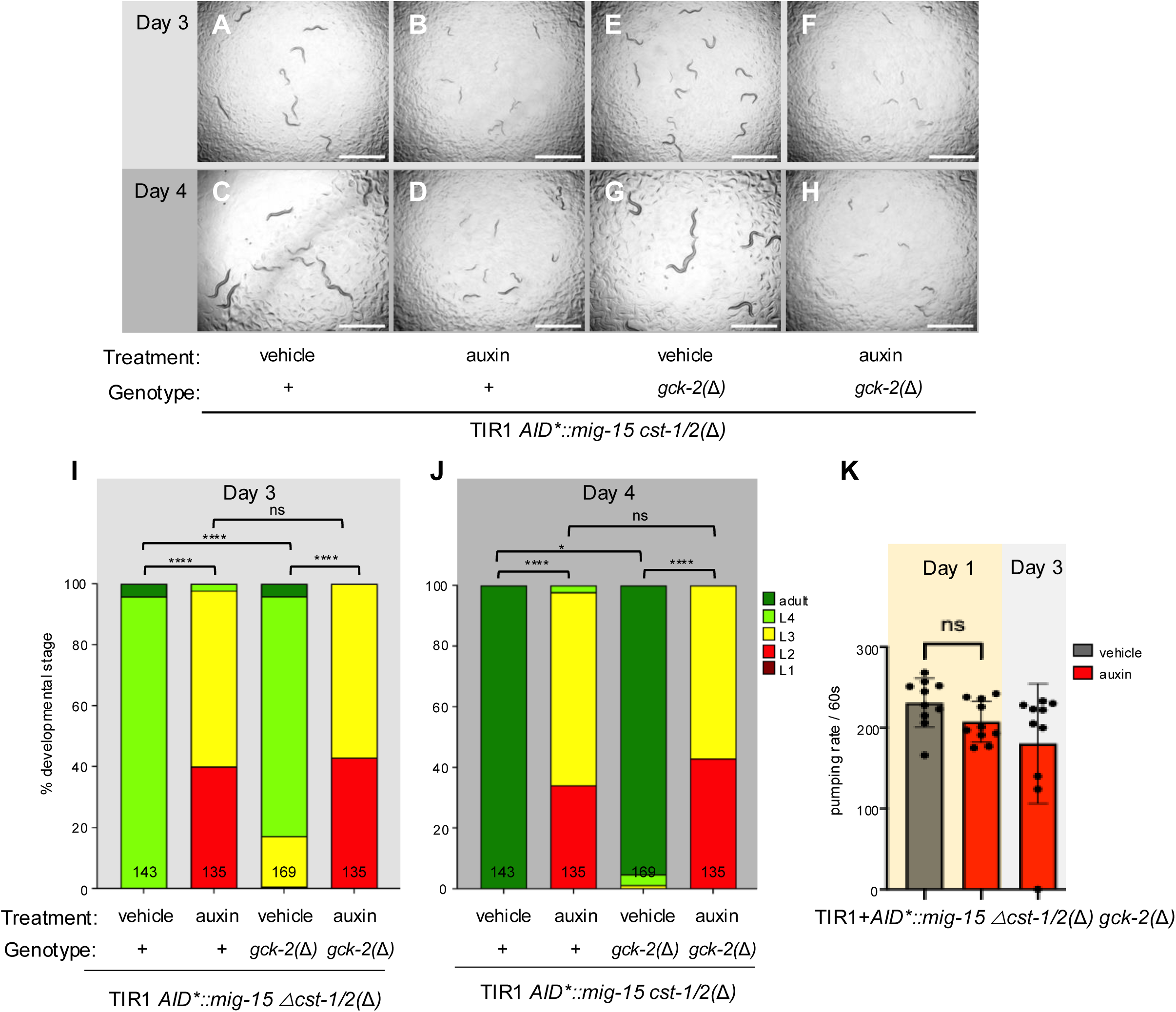
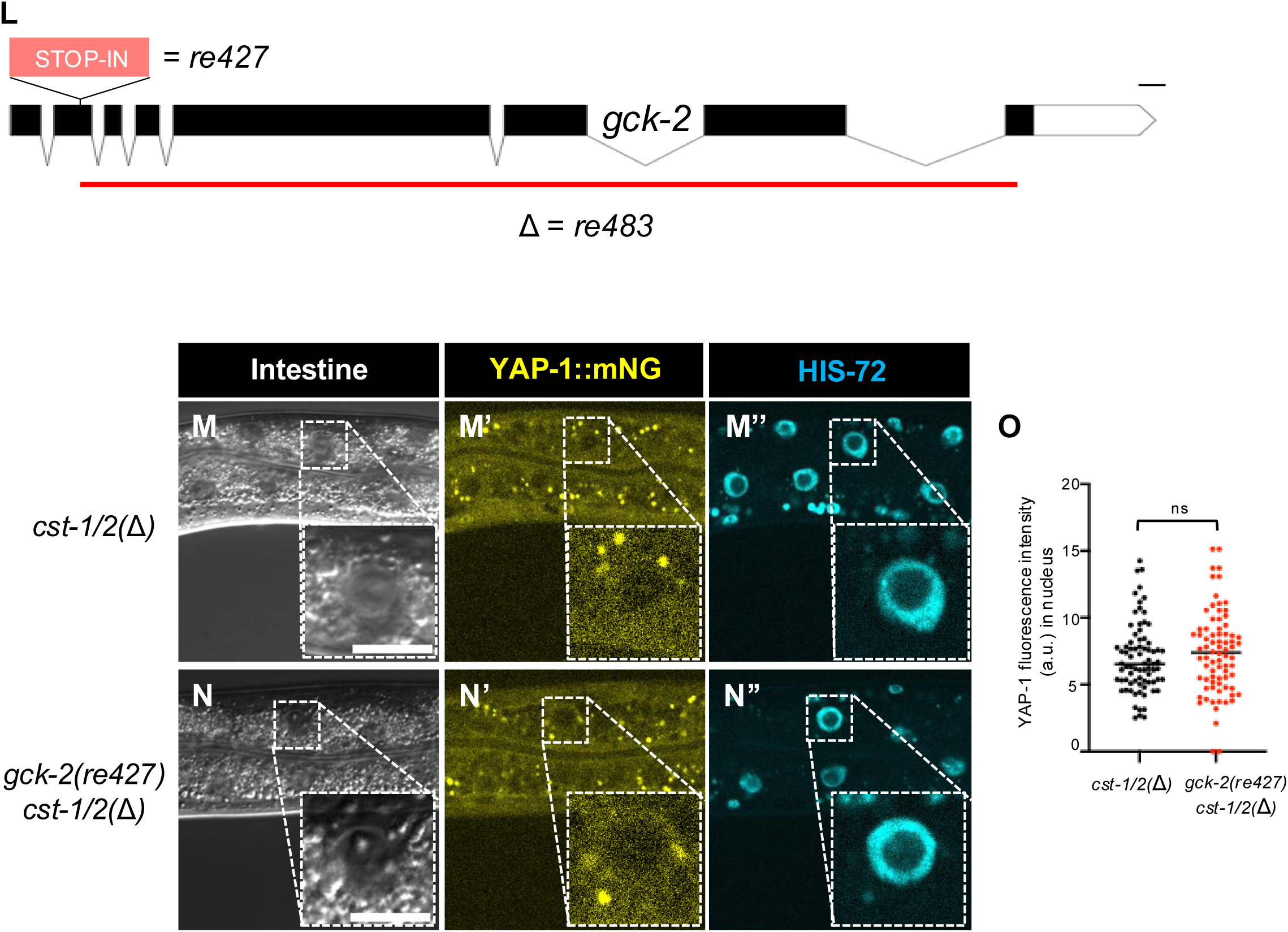

**Figure S6.**
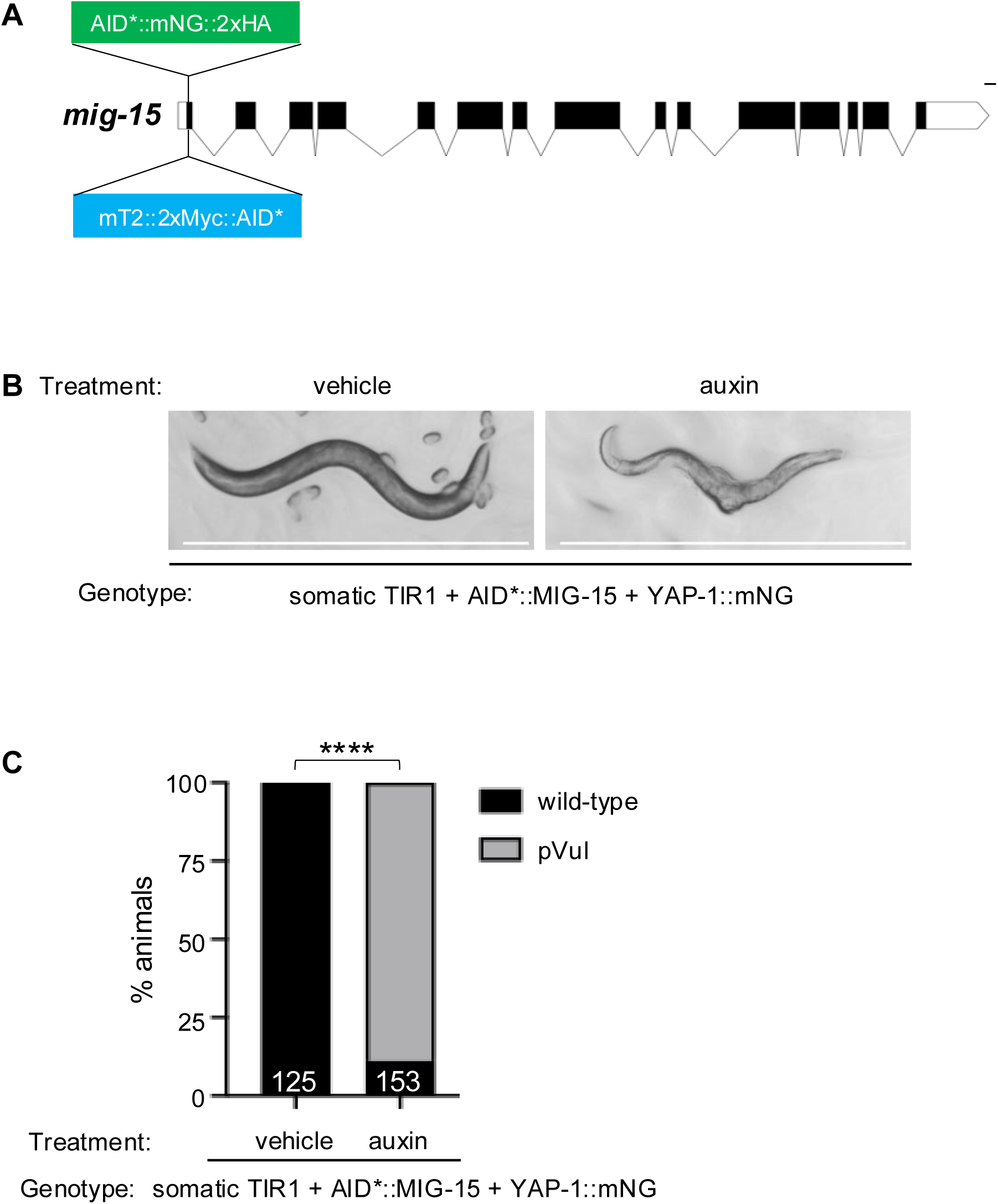

## SUPPLEMENTARY TABLES

**Table S1:** Strain.

| Strain | Genotype |
| --- | --- |
| CA1200 | *ieSi57[eft-3p>TIR1::mRuby + Cbr-unc-119(+)] II; unc-119(ed3) III* |
| DV3709 | *ieSi57[eft-3p>TIR1::mRuby + Cbr-unc-119(+)] II; unc-119(ed3) III; mig-15(re264[AID\*::mNG::2xHA::mig-15]) X* |
| DV3948 | *ieSi57[eft-3p>TIR1::mRuby + Cbr-unc-119(+)] II; unc-119(ed3) III; mig-15(re264[AID\*::mNG::2xHA::mig-15]) cst-1/2(reDf1[re484]) X* |
| DV3969 | *ieSi57[eft-3p>TIR1::mRuby + Cbr-unc-119(+)] II; unc-119(ed3) III; gck-2(re483) V; mig-15(re264[AID\*::mNG::2xHA::mig-15]) cst-1/2(reDf1[re484]) X* |
| DV3611 | *yap-1(re269[yap-1::mNG::2xflag]) X* |
| DV4205 | *ieSi57[eft-3p>TIR1::mRuby + Cbr-unc-119(+)] II; unc-119(ed3) III; yap-1(re269[yap-1::mNG::2xflag]) X* |
| DV4130 | *rrf-3(re390[STOP-IN\*]) II; his-72 (erb77[his-72::linker::mTurquoise2]) III, yap-1(re269[yap-1::mNG::2xflag]) X* |
| DV4186 | *his-72(erb77[his-72::linker::mTurquoise2]) III, yap-1(re269[yap-1::mNG::2xflag]) cst-1/2(reDf4[re452]) X* |
| DV4217 | *wts-1(tm4081)/tmC18[dpy-5(tmIs1236[myo-2p>mCherry])] I* |
| DV4214 | *wts-1(re419[mTurquoise2::2xMyc::AID\*::wts-1]) I; ieSi57[eft-3p>TIR1::mRuby + Cbr-unc-119(+)] II; yap-1(re269[yap-1::mNG::2xflag]) X* |
| DV4288 | *his-72(erb77[his-72::linker::mTurquoise2]) III; gck-2(re427[STOP-IN\*]) V; yap-1(re269[yap-1::mNG::2xflag]) cst-1/2(reDf4[re452]) X* |
| DV4290 | *wts-1(re419[mTurquoise2::2xMyc::AID\*::wts-1]) I; ieSi57[eft-3p>TIR1::mRuby + Cbr-unc-119(+)] II; unc-119(ed3) III* |
| DV4292 | *wts-1(ok753)/tmC18[dpy-5(tmls1236[myo-2p>mCherry])] I* |
| DV4278 | *wts-1(tm4081) I; yap-1(re269[yap-1::mNG::2xflag]) X* |
| DV4297 | *wts-1(re436[STOP-IN\*])/tmC18[dpy-5(tmls1200[myo-2p>Venus]) I* |
| DV4296 | *ieSi57[eft-3p>TIR1::mRuby + Cbr-unc-119(+)] II; unc-119(ed3) III; mig-15(re435[mTurquoise2::2xmyc::AID\*::mig-15]) yap-1(re269[yap-1::mNG::2xflag]) X* |
| DV4337 | *wts-1(re419[mTurquoise2::2xmyc::AID\*::wts-1]) I; reSi2[col-10p>TIR1::F2A::mTaqBFP2::AID\*::NLS] II* |
| DV4338 | *wts-1(re419[mTurquoise2::2xmyc::AID\*::wts-1]) I; reSi12[ges-1p>TIR1::F2A::mTaqBFP2::AID\*::NLS] II* |
| DV4339 | *wts-1(re419[mTurquoise2::2xmyc::AID\*::wts-1]) I; reSi2[col-10p>TIR1::F2A::mTaqBFP2::AID\*::] II; yap-1(re269[yap-1::mNG::2xflag]) X* |
| DV4340 | *wts-1(re419[mTurquoise2::2xmyc::AID\*::wts-1]) I; reSi12[ges-1p>TIR1::F2A::mTaqBFP2::AID\*::NLS] II; yap-1(re269[yap-1::mNG::2xflag]) X* |
| DV4341 | *wts-1(re419[mTurquoise2::2xmyc::AID\*::wts-1]) I; ieSi68[sun-1p>TIR1::mRuby + Cbr-unc-119(+)] II* |
| DV4384 | *ieSi57[eft-3p>TIR1::mRuby + Cbr-unc-119(+)] II; unc-119(ed3) III; mig-15(re435[mTurquoise2::2xmyc::AID\*::mig-15) yap-1(re269[yap-1::mNG::2xflag]) cst-1/2(re479) X* |

**Table S2:**
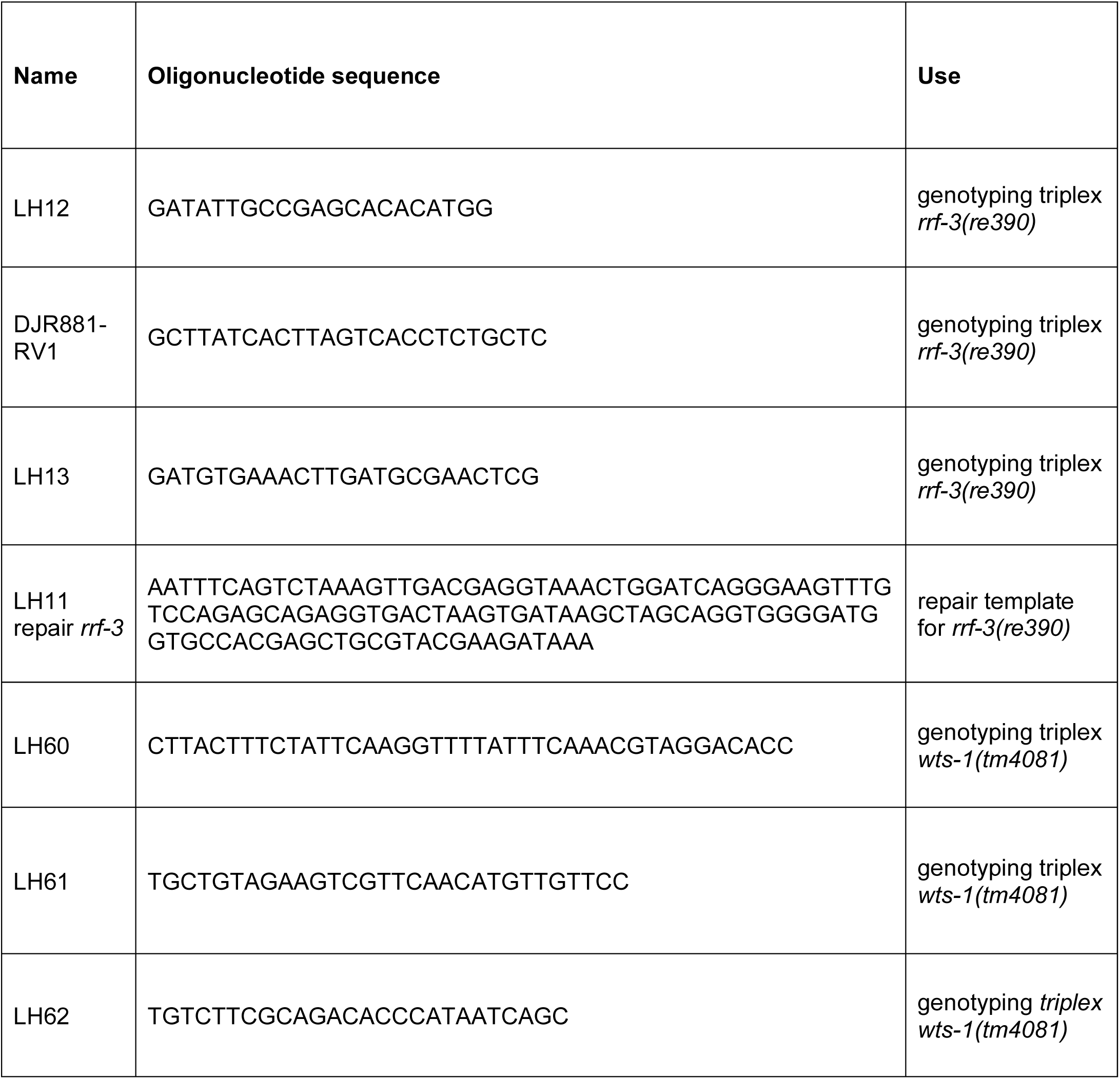
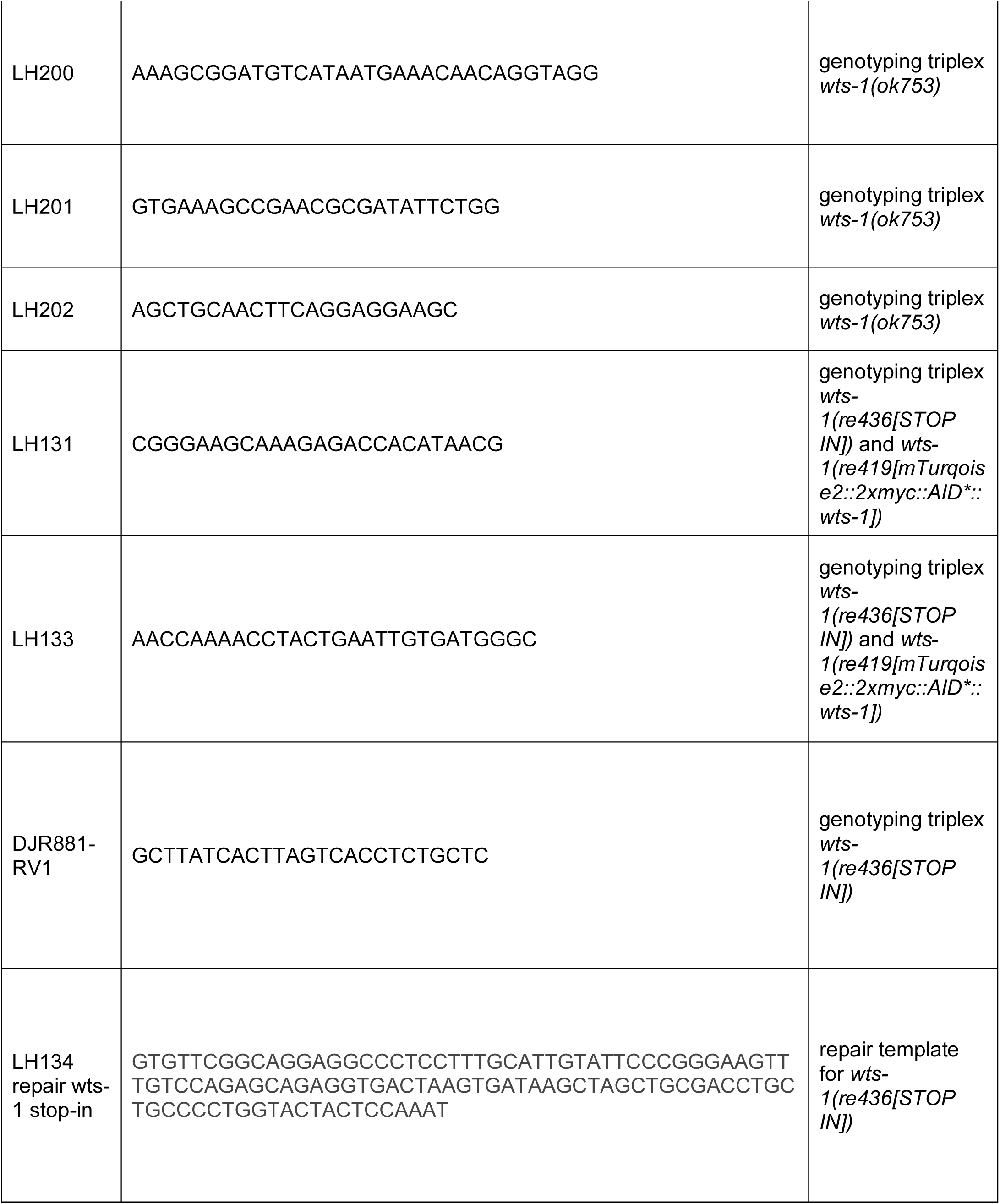
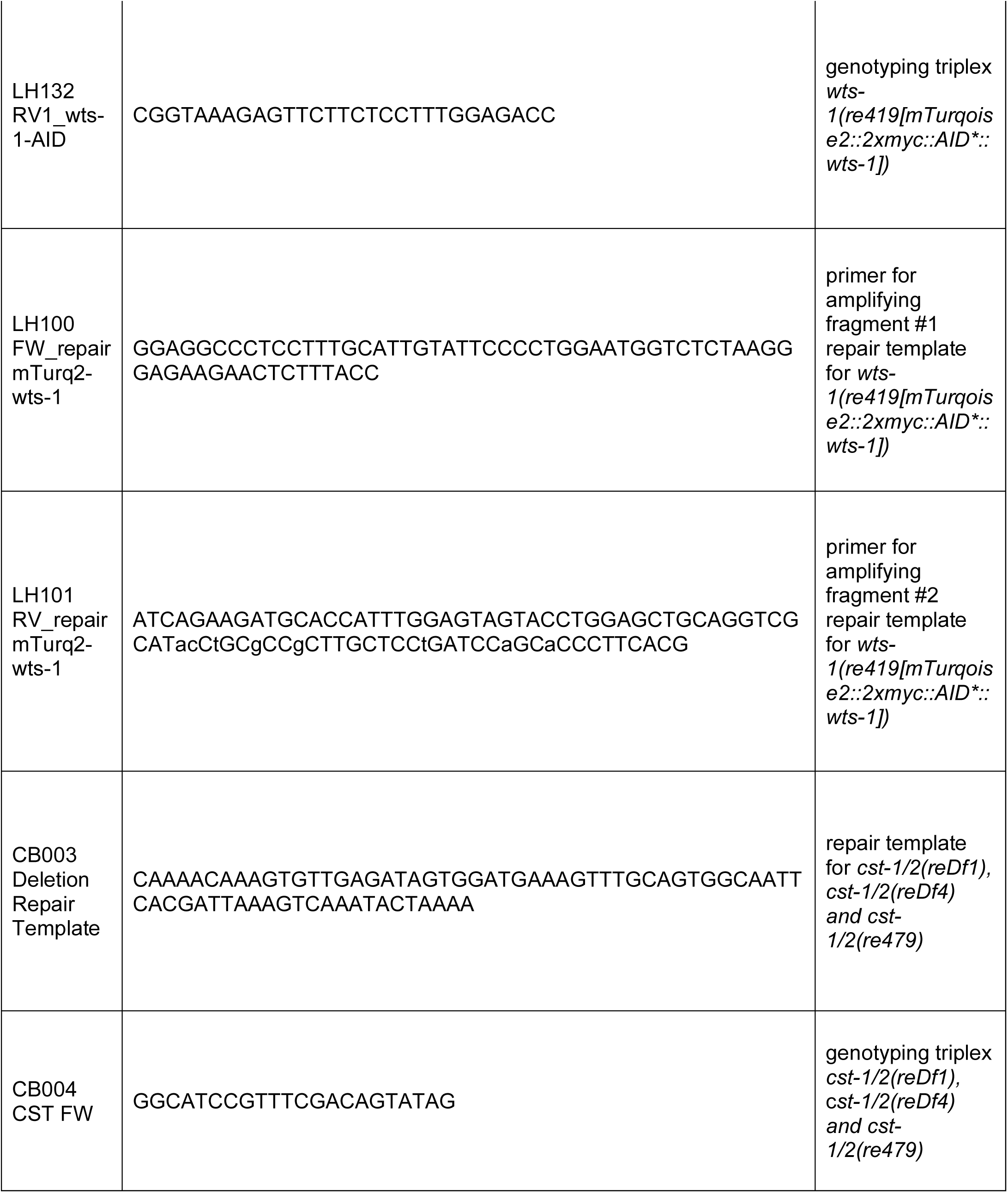
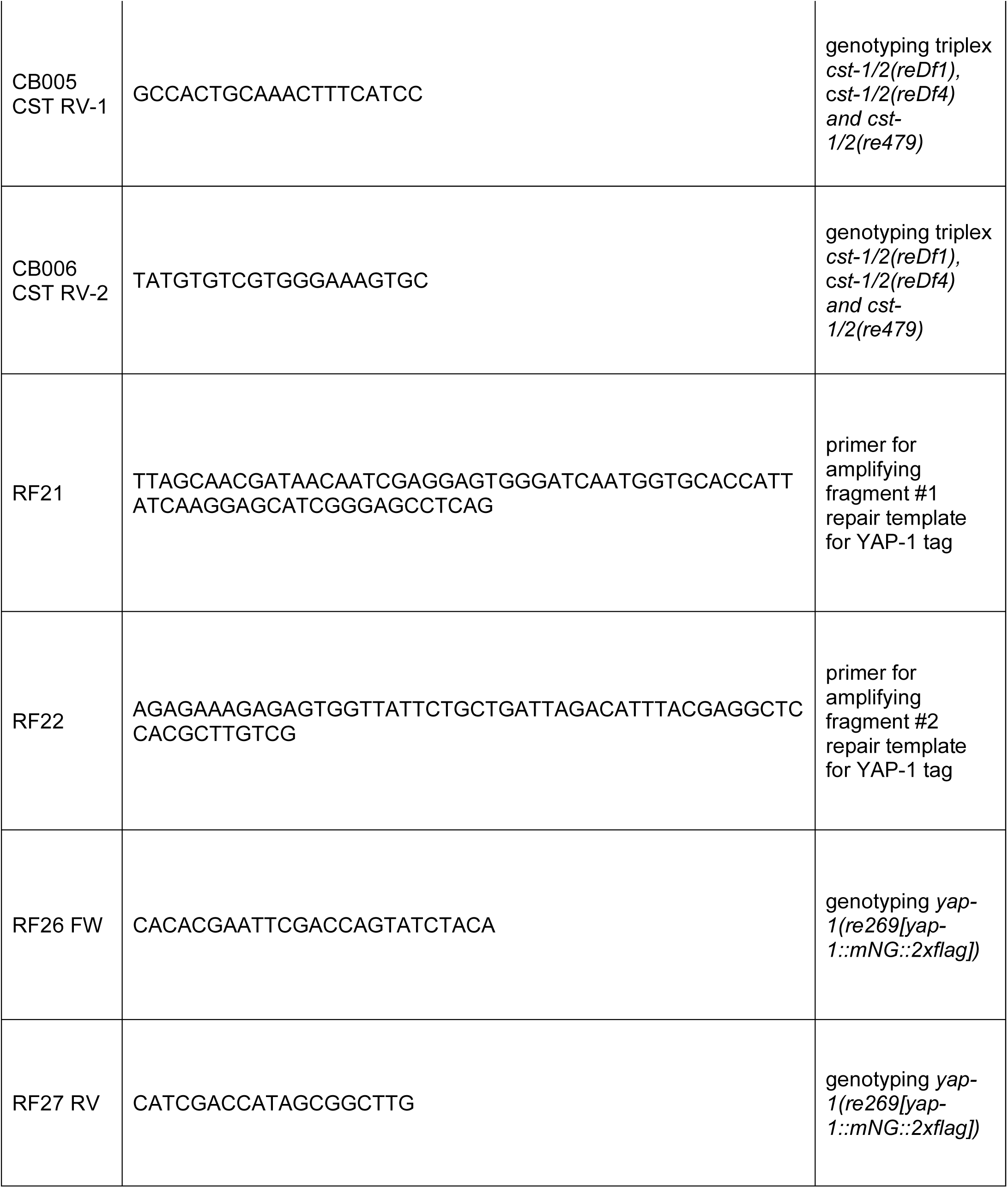
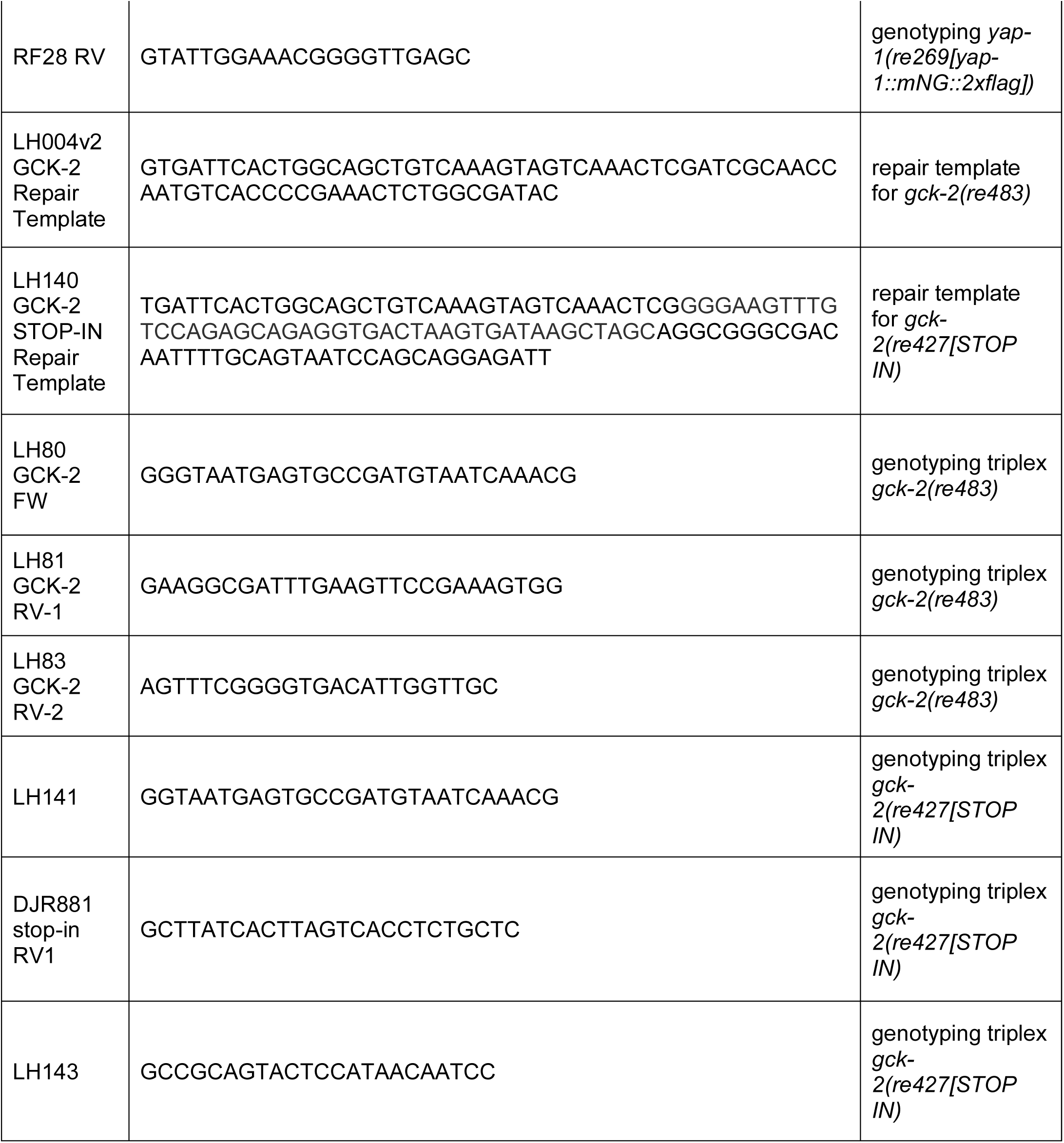
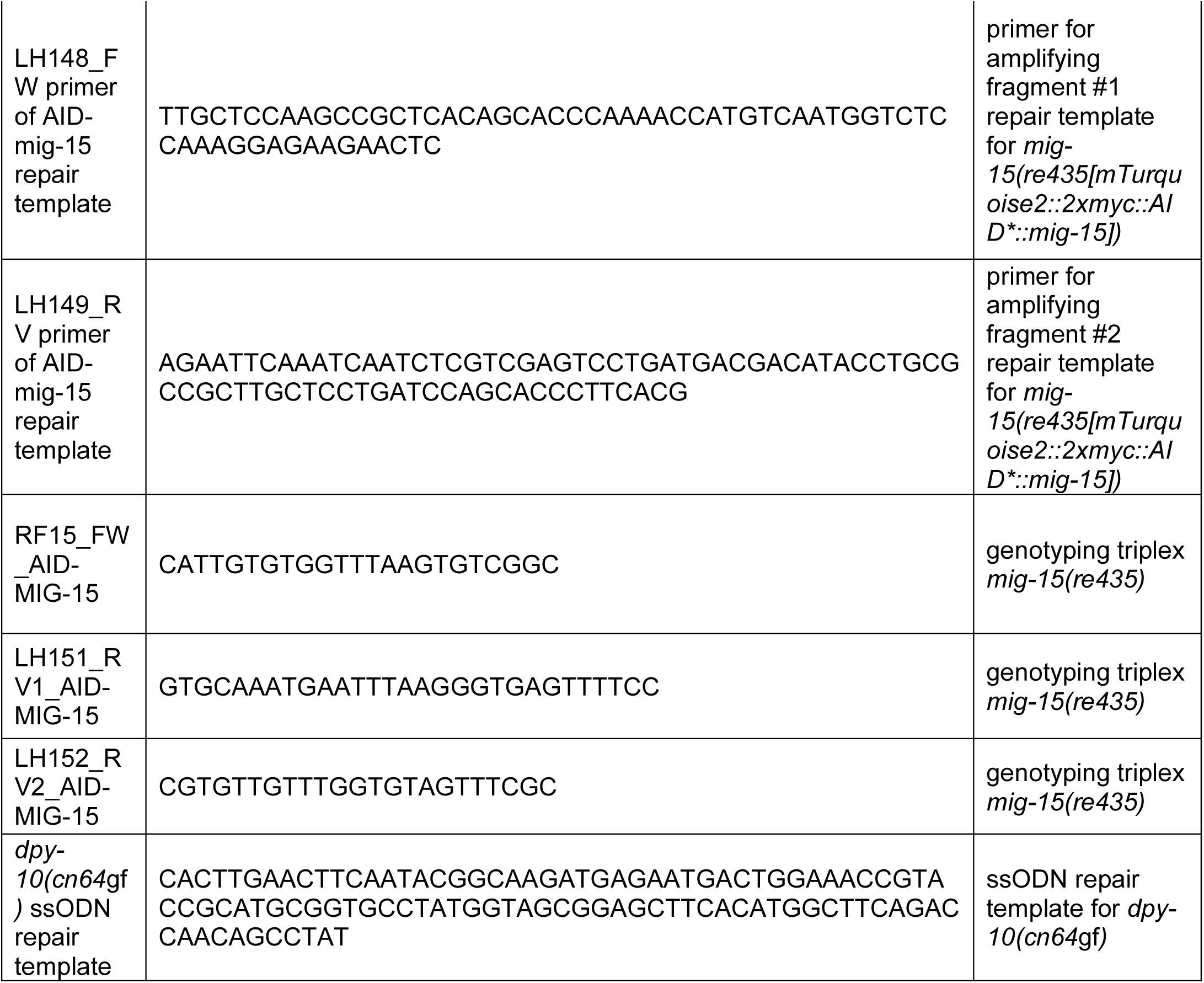
Oligonucleotide.

**Table S3:**
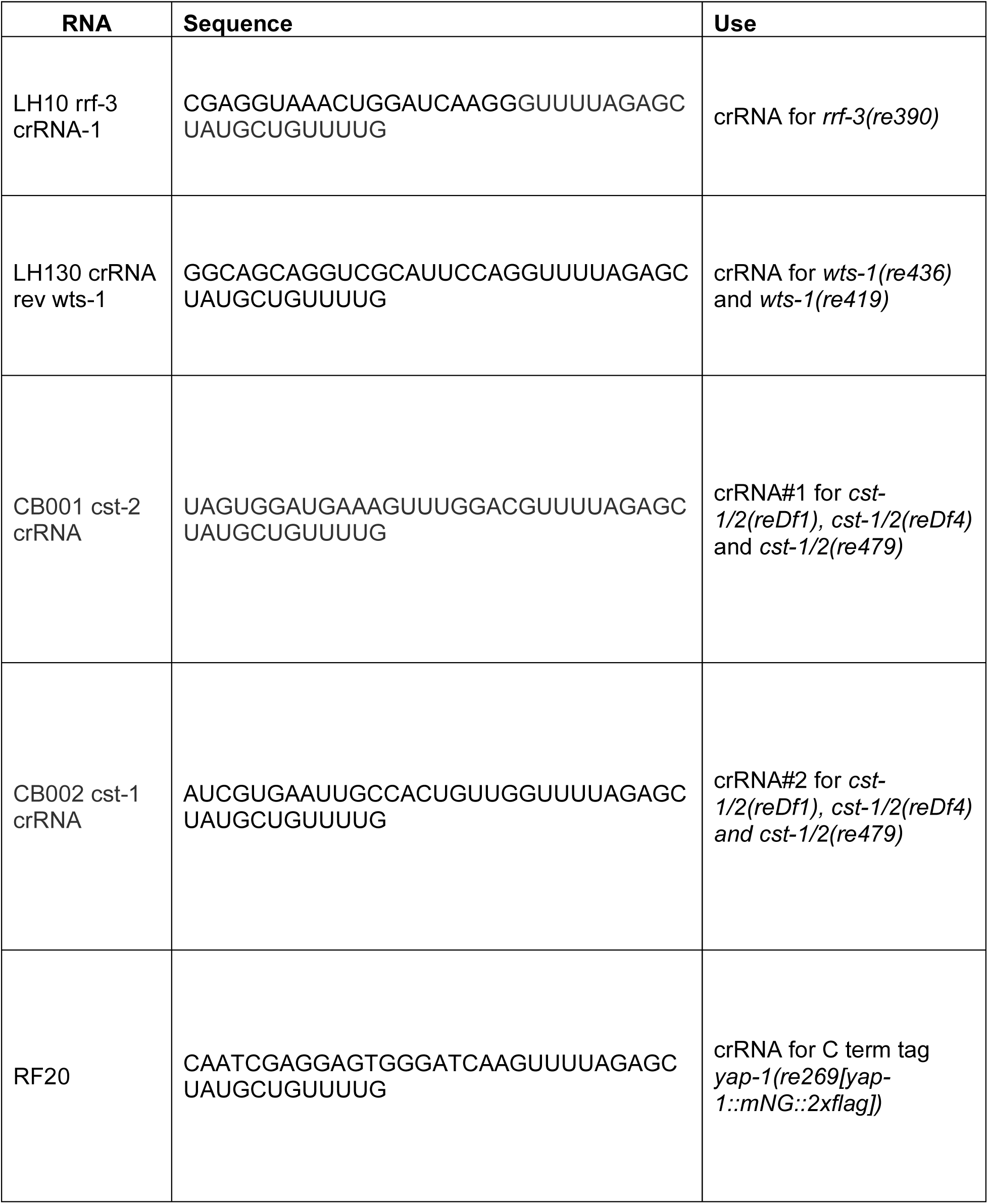
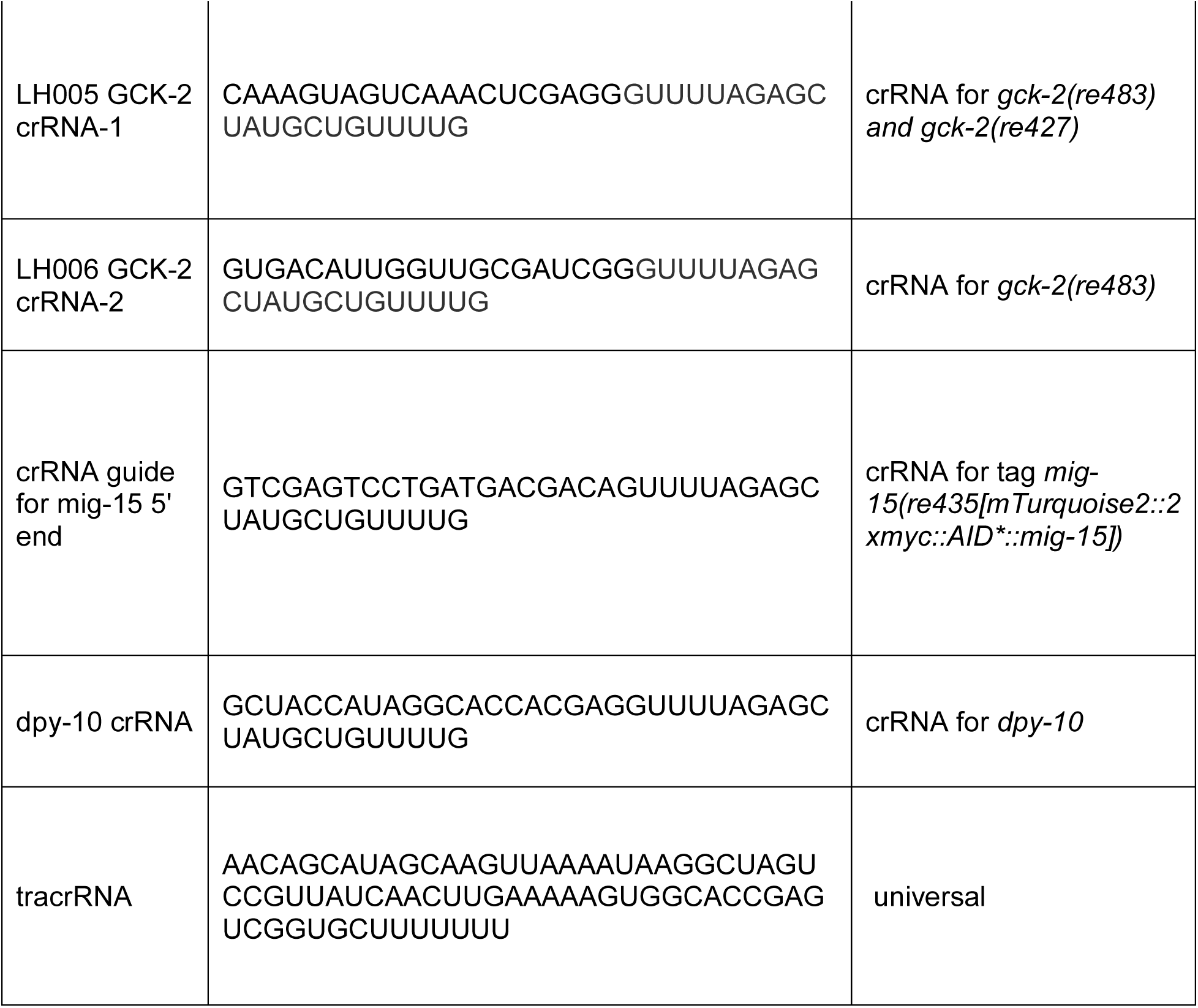
CRISPR guide RNA.

**Table S4:**
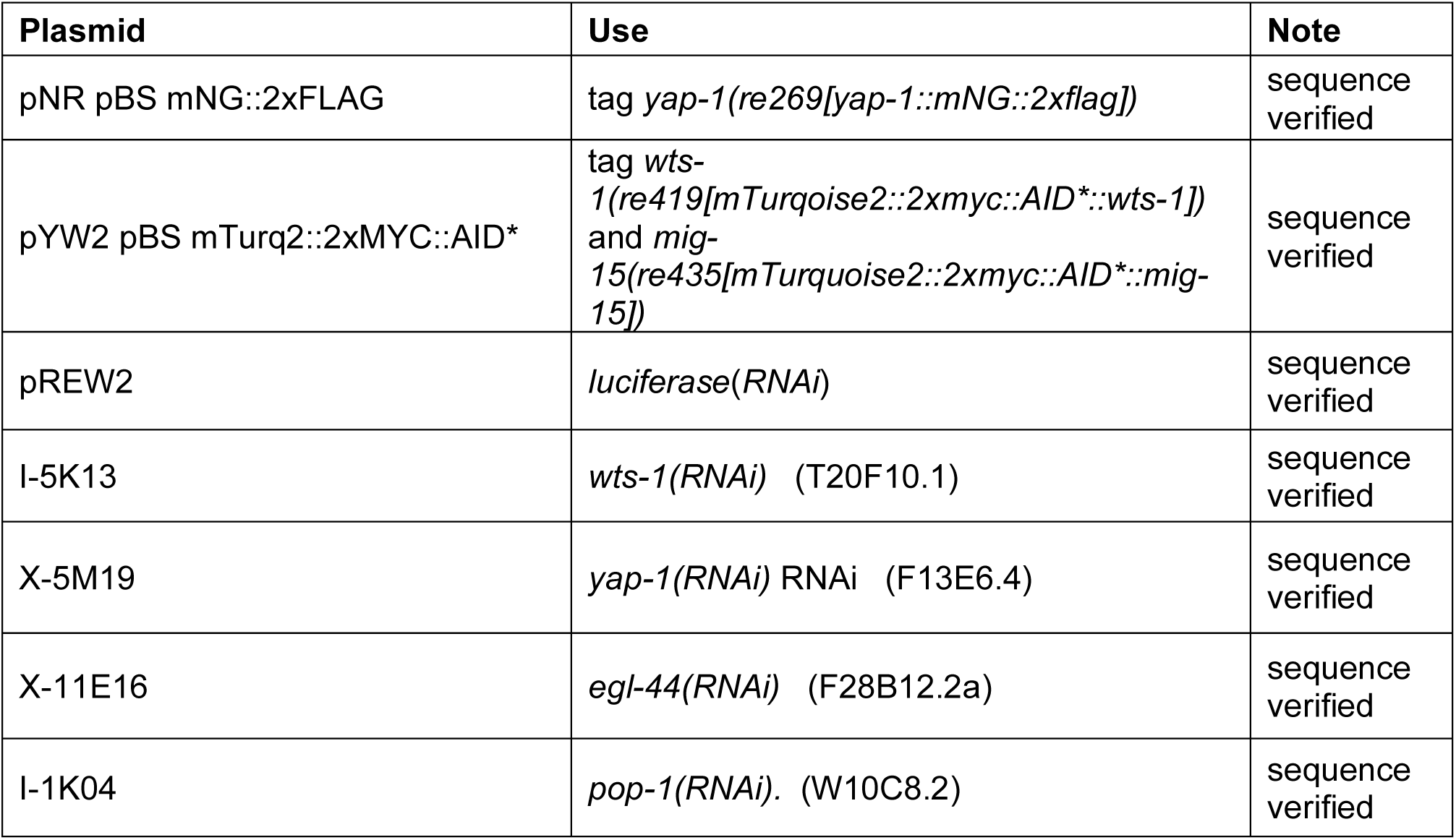
Plasmids.

| **Plasmid** | **Use** | **Note** |
| --- | --- | --- |
| pNR pBS mNG::2xFLAG | tag *yap-1*(*re269*[*yap-1::mNG::2xflag*]) | sequence verified |
| pYW2 pBS mTurq2::2xMYC::AID* | tag *wt*s-1(*re419*[*mTurquoise2::2xmyc::AID\*::wt*s-1])<br>and *mig*-<br>15(*re435*[*mTurquoise2::2xmyc::AID\*::mig*-<br>15]) | sequence verified |
| pREW2 | *luciferase*(*RNAi*) | sequence verified |
| I-5K13 | *wt*s-1(*RNAi*) (T20F10.1) | sequence verified |
| X-5M19 | *yap-1*(*RNAi*) *RNAi* (F13E6.4) | sequence verified |
| X-11E16 | *egl-44*(*RNAi*) (F28B12.2a) | sequence verified |
| I-1K04 | *pop-1*(*RNAi*). (W10C8.2) | sequence verified |

